# “RIG-I mediated neuron-specific IFN type 1 signaling in FUS-ALS induces neurodegeneration and offers new biomarker-driven individualized treatment options for (FUS-)ALS.”

**DOI:** 10.1101/2024.12.02.626340

**Authors:** Marcel Naumann, Stefanie Kretschmer, Banaja Dash, Kevin Peikert, Hannes Glaß, Dajana Großmann, René Günther, Susanne Petri, Annekathrin Rödiger, David Brenner, Francisco Pan-Montojo, Eleonora Aronica, Markus Kipp, Vitaly Zimyanin, Jared Sterneckert, Torsten Grehl, Tobias M. Böckers, Patrick Oeckl, Min Ae Lee-Kirsch, Andreas Hermann

**Affiliations:** Translational Neurodegeneration Section “Albrecht Kossel”, Department of Neurology, University Medical Center Rostock, University of Rostock, Rostock, Germany; Department of Pediatrics, Medizinische Fakultät Carl Gustav Carus, Technische Universität Dresden, Dresden, Germany; Department of Neurology, Technische Universität Dresden, Dresden, Germany; Deutsches Zentrum für Neurodegenerative Erkrankungen (DZNE) Dresden, Dresden, Germany; Department of Neurology, University Hospital Carl Gustav Carus at Technische Universität Dresden, Dresden, Germany; Deutsches Zentrum für Neurodegenerative Erkrankungen (DZNE) Dresden, Dresden, Germany; Department of Neurology, Hannover Medical School, Hannover, Germany; Department of Neurology, Jena University Hospital, Jena, Germany; University Hospital Ulm, Department of Neurology, Ulm, Germany; Dept. of Psychiatry and Psychotherapy at the Klinikum LMU, Munich, Germany and Neurologische Klinik Sorpesee, Sundern, Germany; Amsterdam UMC, University of Amsterdam, Department of (Neuro)Pathology, Amsterdam Neuroscience, Amsterdam, The Netherlands; Rostock University Medical Center, Institute of Anatomy, Rostock, Germany; Department of Molecular Physiology and Biological Physics, School of Medicine, University of Virginia, Charlottesville, VA, 22903, USA; (S.R.) Center for Membrane and Cell Physiology, School of Medicine, University of Virginia, Charlottesville, VA, 22903, USA; Center for Regenerative Therapies TU Dresden (CRTD) and the Medizinische Fakultät Carl Gustav Carus, Technische Universität Dresden, Dresden, Germany; Department of Neurology, Alfried Krupp Hospital, Essen, Germany; Institute of Anatomy and Cell Biology, University of Ulm, Ulm, Germany; German Center for Neurodegenerative Diseases (DZNE) Ulm, Ulm, Germany; University Centre for Rare Diseases, University Hospital Carl Gustav Carus, Technische Universität Dresden, Dresden, Germany; German Center for Child and Adolescent Health (DZKJ), partner site Leipzig/Dresden, Dresden, Germany; German Center for Neurodegenerative Diseases (DZNE) Rostock/Greifswald, Rostock, Germany; Center for Transdisciplinary Neurosciences Rostock (CTNR), University Medical Center Rostock, University of Rostock, Rostock, Germany

**Keywords:** pathogen-associated molecular patterns (PAMPs), RIG-I like receptors, double-stranded RNA, cGAS-STING pathway, interferon type 1, RNA-sequencing, RIG-I

## Abstract

Recent research has demonstrated significant aberrant activation of the innate immune system in ALS model systems due to mutations in SOD1, TARDBP and C9orf72 through stimulation of the TBK1-IRF3 pathway. This pathway can be activated, for example, by cGAS-STING-dependent sensing of cytosolic DNA that accumulates as a result of chronic DNA damage and defective mitochondria, both of which have been identified as early pathology in FUS-ALS spinal motor neurons (sMNs). Therefore, we analysed innate immune pathways in isogenic and non-isogenic FUS^mut^ iPSC-derived sMNs, which revealed upregulation of interferon-stimulated genes (ISGs) and activation of the TBK1-IRF3 pathway in FUS^mut^ sMNs. Notably, we found evidence for accumulation of cytosolic dsRNA and its sensor RIG-I in FUS-ALS. RIG-I, but not MDA5, was found to be significantly upregulated in FUS^mut^ sMNs, and siRNA-mediated knockdown abolished the increased IFN1 activation in FUS^mut^ sMNs. In post-mortem analysis, RIG-I was highly expressed in the remaining α-MNs. IFN treatment of FUS^wt^ sMNs phenocopied the axonal degeneration of FUS^mut^ sMNs. Mechanistically, DNA damage induction did not increase ISG expression, but dsRNA was increased in the mitochondria of FUS^mut^ sMNs. Mitochondrial transcription, a known source of dsRNA, was found to be upregulated in compartmental axonal RNAseq analysis and its inhibition reduced ISGs in FUS-ALS sMNs. Furthermore, the JAK-STAT inhibitor ruxolitinib alleviated the upregulated ISG expression and reversed the axonal degeneration of sMNs. Finally, we analysed ISG expression in peripheral blood samples from 18 FUS-ALS patients, eight of whom had a significantly elevated interferon signature. Blood ISGs correlated with disease progression rate and negatively with disease duration. RIG-I-mediated innate immune activation in sMNs may be an interesting novel individualised biomarker-driven therapeutic target in (FUS-) ALS.

**A one-sentence summary of your paper:** RIG-I-I-mediated innate immune activation is found in FUS-ALS spinal motor neurons caused by cytosolic dsRNA accumulation due to mitochondrial transcriptional activation and is amenable to JAK-STAT inhibition and might thus be an interesting novel individualized biomarker-driven therapeutic approach in (FUS-) ALS

## Introduction

Despite decades of intensive research, our understanding of the specific pathomechanisms of selective motor neuron degeneration in amyotrophic lateral sclerosis (ALS), a devastating neurodegenerative disease, is incomplete. In addition to perturbations in proteostasis, autophagy, and endoplasmic reticulum (ER) stress, among others, (for review see ^1^), there is growing evidence that innate immune pathways promote sterile neuroinflammation, which may contribute significantly to the pathomechanisms of ALS and other neurodegenerative diseases. ^2,3^. In detail, mitochondrial distress and increased production of reactive oxygen species fatally synergize to induce nuclear or mitochondrial DNA damage and increase the permeability of the mitochondrial transition pores ^4^. This leads to the translocation of damaged nuclear and/or mitochondrial DNA into the cytosol, which is recognized as “foreign” by conserved cellular mechanisms whose actual purpose is to elicit a response to get rid of pathogens after sensing their foreign DNA during infection. In neurodegenerative diseases, however, these mechanisms become suicidal. Firstly, recognition of cytosolic double-stranded DNA is mediated by cyclic guanosine monophosphate–adenosine monophosphate (cGAMP) synthase (cGAS), which activates the adaptor protein stimulator of interferon genes (STING) ^5^. Secondly, double-stranded RNA derived from damaged mitochondria is sensed by members of the Retinoic acid-inducible gene I (RIG-I)-like receptor (RLR) family, such as RIG-I (also known as DDX58), which interacts with the mitochondrial antiviral signaling protein (MAVS) ^6^. Both adaptor proteins, STING and MAVS, recruit TANK-binding kinase 1 (TBK1) and are phosphorylated by it at a conserved c-terminal consensus motif ^7^, which is independent of other cellular functions of TBK1, such as in autophagy. In particular, this allows the recruitment and phosphorylation of interferon regulatory factor 3 (IRF3). Phosphorylated IRF3, in turn, dimerises and shuttles to the nucleus to induce the expression of interferon-stimulated genes (ISGs), resulting in the activation of interferon type 1 (IFN-1), stimulation of the TNF-alpha pathway and the upregulation of several other inflammatory mediators, pro-apoptotic genes and chemokines. In unaffected tissues, there is a low or ’tonic’ production of IFN-1, which is essential for normal cellular function and host defence against microbial pathogens. This physiological homeostasis can be thrown out of balance by TBK1-IRF3 signaling in, for example, senescent microglia, putting them in a reactive state that causes non-cell autonomous neuronal neurotoxicity ^2^.

Importantly, C9orf72 mutations have been shown to cause over-activation of the innate immune system in mouse models, cell culture systems and patient blood samples ^8–10^. This has been experimentally attributed to activation of the TBK1-IRF3 pathway. However, different primary mechanisms for promoting this pathway have been described: On the one hand, it was shown that loss of C9orf72 led to reduced clearance of STING by autophagy, resulting in chronic TBK1 stimulation ^10^. On the other hand, bidirectional transcription of C9orf72 repeats was shown to generate sense and antisense mRNA with subsequent dsRNA formation. This was shown to trigger MDA5 activation leading to neuronal loss in C9orf72 HRE mutant mouse model systems and human iPSC-derived sMNs ^9^. In TARDBP mutant model systems, STING activation was identified as a result of mitochondrial DNA release due to TDP-43-mediated opening of the mitochondrial permeability transition pore (mPTP) ^4^. In a SOD1 mutant ALS mouse model, mitochondrial misfolded SOD1 was found to promote mtDNA and RNA:DNA release into the cytosol independently of mPTP, resulting in STING activation ^11^. Finally, recent evidence from human post-mortem studies suggests an elemental, cell-autonomous abundance of STING in cortical and spinal motoneurons in sporadic and some familial forms of ALS ^12^.

Overall, overactivity of the cGAS-STING/RIG-I and downstream TBK1-IRF3 signaling pathways has recently been identified as a major contributor to neurodegeneration in genetic models of ALS. However, it remains unclear whether similar mechanisms are involved in FUS-ALS. Previously, we and others have shown that FUS-ALS spinal motor neurons (sMNs) accumulate DNA damage and depolarised axonal mitochondria ^13–15^. Therefore, the IFN-1 response may also be central to the pathogenesis of FUS ALS. There is considerable evidence that DNA damage may precede (axonal) neurodegeneration in several ALS models, including FUS ^13,16^. However, the relationship between the two remains to be elucidated. Some reports suggest that TNFα treatment induces axonal dysfunction ^17^ and imply a neurotoxic effect of chronic IFN1 exposure on neurons ^18^. Therefore, we aimed to investigate the role of IFN1 signaling in FUS-ALS to determine if there is a neuronal, cell-autonomous activation of IFN1 signaling and, if so, through which specific pathway it is mediated and whether this IFN1 signaling might be a link between DNA damage and the observed axon degeneration. The canonical IFN1 pathway activates the Janus kinase (JAK) signal transducer and activator of transcription (STAT) pathway. Since JAK-STAT inhibitors are already FDA-approved for several non-neurological diseases, we wanted to further investigate whether they could alleviate neurodegeneration in ALS and thus represent an interesting repurposing strategy for a biomarker-driven FUS-ALS therapeutic approach.

## Results

### Activation of interferon type 1 response in iPSC-derived FUS^mut^ mutant motor neurons

Previously, we have shown that FUS-ALS sMNs accumulate DNA damage and depolarised axonal mitochondria ^13,14^. This led us to hypothesise that there may be at least one mechanism that triggers an innate immune response, either through mitochondrial or nuclear DNA leakage. To address whether FUS mutations lead to innate immune activation within the neurons themselves, we assessed IFN1 pathway stimulation by measuring a set of interferon-stimulated genes (ISGs) (Fig. 1A+B) by RT-qPCR under baseline conditions. This was performed in iPSC-derived sMNs cultures devoid of glial cells, comparing the FUS-P525L mutation with an isogenic control (Fig. 1A). These cell lines were previously edited by CRISPR-Cas9n to create an isogenic condition and add GFP to the C terminus of FUS in both cell lines ^13^. Three additional (patient-derived) FUS mutant iPSC-derived sMNs cell lines (R521C, R521L, R495Qfs*527) were analysed for ISG expression, which was normalised to two non-isogenic FUS^WT^ control sMNs lines (Figure 1B). All lines were previously established and published, including negative genetic testing for C9ORF72, FUS, TARDBP and SOD1 for the WT cell lines ^13^. From the latter study, we knew that significant mitochondrial phenotypes and DNA damage accumulation were not present until at least 3 weeks during neuronal maturation, which is why the ISG measurement was performed at this time point. For FUS-P525L sMNs, there was a robust upregulation for all ISGs examined (Fig. 1A). For the non-isogenic lines, there was still a significant upregulation of some ISGs (Fig. 1B).

**Figure 1.**
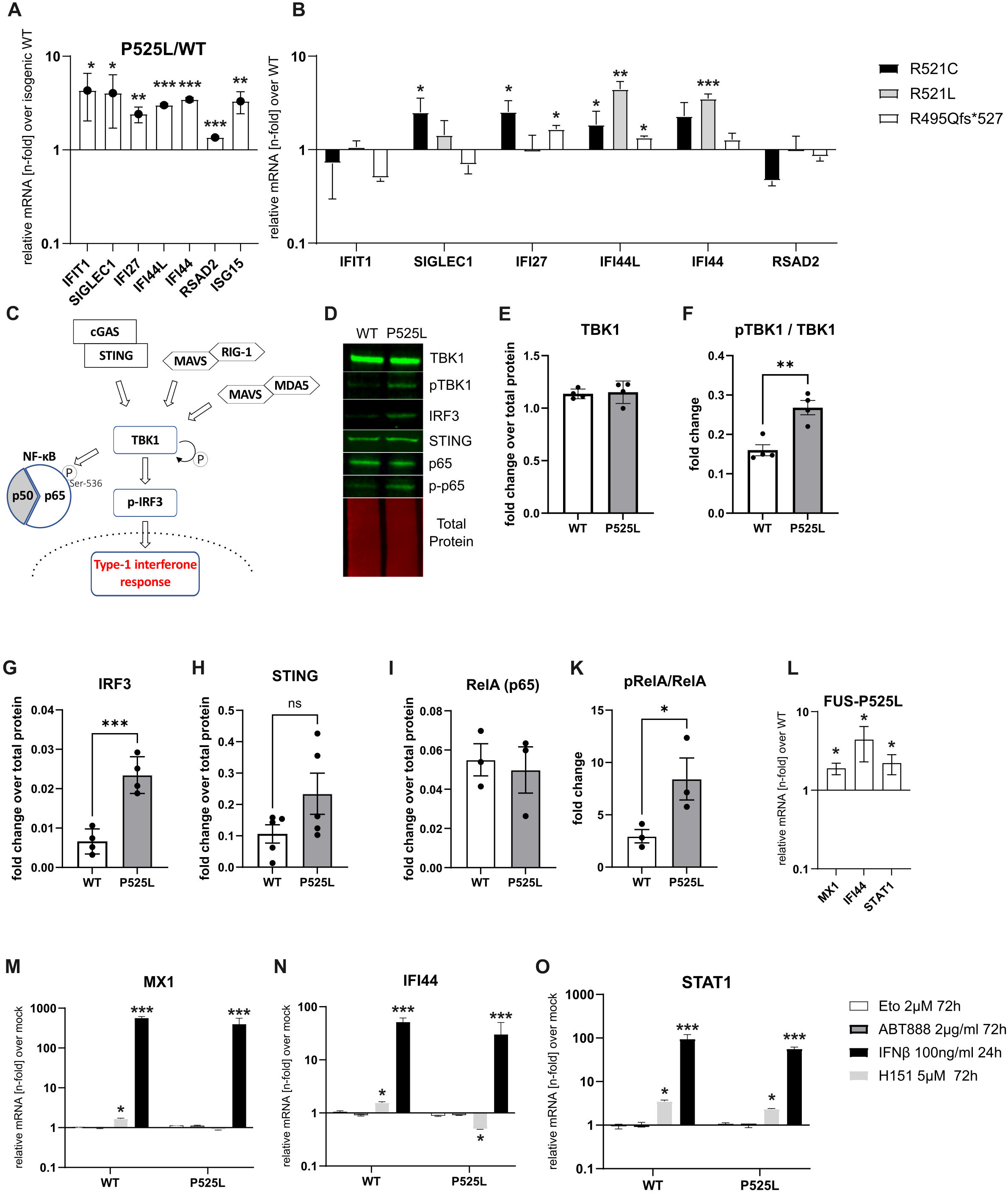
Innate immune activation in iPSC-derived spinal motoneurons with mutations in *FUS*. (A, B) RT-qPCR for a set of ISGs, as indicated on the x-axis in sMNs with a FUS-P525L mutation, normalized to an isogenic control line (A) and three other sMNs lines with distinct C-terminal *FUS* mutations normalized to two non-isogenic FUS^wt^ sMNs lines (B), unpaired t-test. (C) sketch of the TBK1-IRF3 pathway. TBK1 activation by either RIG-I or cGAS sensing results in downstream transcriptional activation of a type-1 IFN response following the translocation of phosphorylated IRF3 into the nucleus or phosphorylation of p65 at Ser-536. (D) Example western blot scan indicating protein expression of markers of the TBK1-IRF3 pathway in WT and isogenic FUS-P525L sMNs. (E-H) Quantification of (D) normalized to a total protein staining, n=4, unpaired t-test. (I,K) Quantification of RelA and p-RelA WB band intensity, n=3, unpaired t-test. (K) (L) RT-qPCR for selectively IFN-1 dependent ISGs (MX1, IFI44, and STAT1) in FUS-P525L sMNs normalized to isogenic control and (M-O) influence of different treatment strategies on their expression normalized to mock treatment, one-way ANOVA, Tukey’s post hoc. DNA-damaging approaches (etoposide, PARP inhibitor ABT888) did not change their expression in FUS^mut^ or FUS^wt^. Similarly, the STING inhibitor H151 did not lower the ISG expression in mutant sMNs. Treatment with IFN-beta as a positive control.

We then tested for activation of the TBK1-IRF3 pathway, based on the hypothesis that this pathway might be responsible for the observed ISG stimulation. As shown in the sketch in Figure 1C, the TBK1-IRF3 pathway acts as a common trunk of several upstream signaling pathways that function in sensing different pathogen-associated molecular patterns (PAMPs). We further focused on the isogenic P525L and corresponding WT sMNs lines as they showed the most homogeneous IFN1 pathway activation at the mRNA level. As expected, there was no difference in the baseline level of TBK1 in both FUS^mut^ and FUS^WT^ sMNs (Fig. 1E). However, we found a significant increase in Ser172 TBK1 phosphorylation (i.e. activation of TBK1) in mutant sMNs (Fig. 1F). There was also a significantly higher signal for IRF3 in mutant neurons (Fig. 1G), together suggesting pathway activation. In addition, we found significantly increased p65 phosphorylation, consistent with NFκB activation (Fig. 1H-K).

### DNA damage induction did not induce interferon type 1 response in iPSC-derived FUS mutant motor neurons

We and others have shown that the role of FUS in the DNA damage response is downstream and dependent on poly (ADP-)ribose polymerase 1 activation ^13,19^. To investigate whether DNA damage induction is a major source of ISG up-regulation in our model, we treated cells for 3 days with 2 µM etoposide - a topoisomerase inhibitor known to induce DNA damage - or 2 µg/ml of the PARP inhibitor ABT888. For this experiment, we assessed the expression of ISGs (MX1, STAT1 and IFI44), which were significantly upregulated in FUS-P525 sMNs (Fig. 1L). Unexpectedly, DNA-damaging treatments did not lead to an upregulation of these ISGs in either cell line (Fig. 1M-O) although we knew from previous studies that this experimental design led to a robust DNA damage induction in identical experimental settings ^13^, as indicated by an increased number of γH2A.x nuclear foci following etoposide treatment (Suppl. Fig 1A). This suggests that activation of nuclear DNA damage may not be the source of the observed ISG activation.

### Inhibition of STING signaling does not alleviate interferon type 1 activation in iPSC-derived FUS mutant motoneurons

Having found evidence for upregulation of the TBK1 pathway and ISG mRNA, we next sought to identify the upstream mechanism. Notably, we found no significant difference in STING expression (Fig. 1H-K). Next, we asked whether H151, an established small molecule inhibitor of STING by blocking STING palmitoylation ^20^, could reduce the increased ISG activation found in FUS-P525L sMNs, as previously shown in iPSC-derived neurons with different genetic ALS mutations ^12^. However, we did not observe a systematic decrease of ISG levels in the FUS^mut^ sMNs upon exposure of H151. Interferon-beta was used to control for the extent of possible ISG stimulation in this cell type (Fig. 1M-O).

### Cytosolic dsRNA in iPSC-derived FUS^mut^ motoneurons

Alternative pathways for ISG activation include RIG-I/MDA5 sensing of dsRNA species. We therefore hypothesised that there might be an overabundance of cytosolic dsRNA triggering the observed innate immune stimulation in FUS^mut^ sMNs. To address this question, super-resolution AiryScan2 LSM microscopy was performed to identify the cytosolic dsRNA immunofluorescence signal using the well-established J2 antibody, which recognises dsRNA species of at least 40 bp. To separate free cytosolic from mitochondrial dsRNA, we generated masks from the immunofluorescence signal of the mitochondrial marker HSP60. We quantified both the mitochondrial and non-mitochondrial/free-cytosolic compartments of the dsRNA immunofluorescence signal. This was done in FUS-P525L and compared to the isogenic control sMNs (Fig. 2A). To demonstrate the quantification technique, the insets surrounded by the dotted line in Figure 2A show the masks generated by Fiji that recognised the cytosolic dsRNA signal and were used to measure the pixel area.

**Figure 2.**
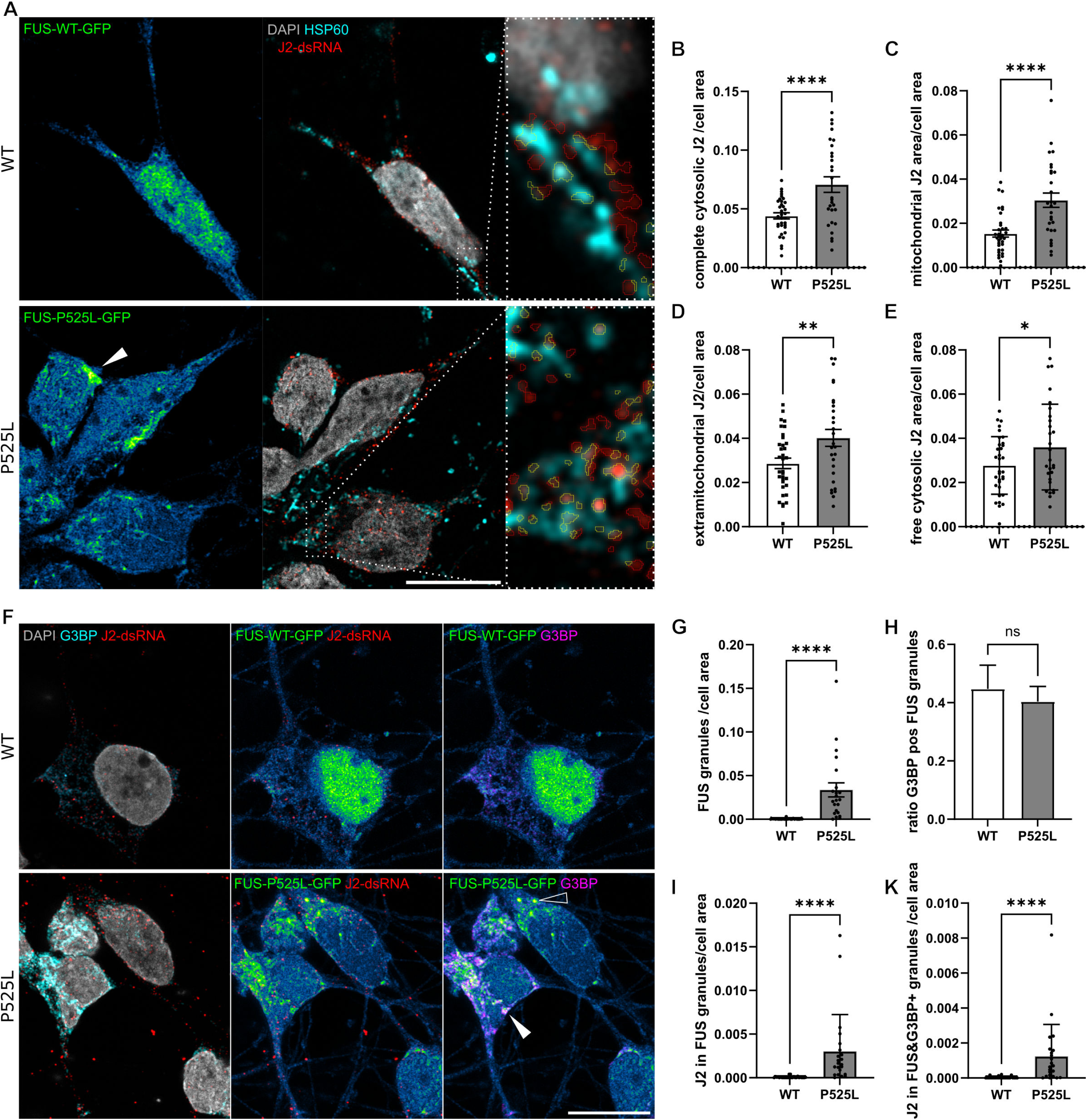
Assessment and quantification of cytosolic dsRNA in relation to mitochondrial compartments in iPSC-derived sMNs. (A) IF image panel of WT and FUS-P525L sMNs, scale bar 10µM. Cells were stained against the mitochondrial matrix protein HSP60 and an anti-dsRNA (J2) antibody (A). Nuclear staining was performed with DAPI. C-terminally tagged FUS-GFP indicates FUS presence in the different conditions and is visualized via the Green-Fire-Blue LUT in Fiji. Note the nuclear loss of FUS-P525L-GFP. Dotted magnification boxes show the individual recognition of dsRNA (A) as region of interest (ROI) areas (yellow line) in Fiji inside or outside of the HSP60 area. (B-E) Quantification of the cytosolic J2-dsRNA ROI area (B) within the HSP60 compartment (C) and outside of it (D, extramitochondrial cytosolic) and outside of HSP60 and FUS granule area (E, free cytosolic). Normalization was as per whole cell area of the cell, respectively. (B-E) Significantly more dsRNA signal per cell was detected in FUS-P525L sMNs compared to control in all conditions (t-test, n=3). (F) IF image panel of WT and FUS-P525L sMNs indicating J2-dsRNA in relation to G3BP as a marker of stress granules, scale bar 10µM. Note the spontaneous formation of complex FUS-GFP granules in FUS-P525L sMNs (see G for quantification), which were partially double-labeled with G3BP positive granules (filled white arrowhead), but not always (arrowhead). Around 40% of all FUS granules was G3BP double-positive without difference between the cell lines (H). (I, K) Enrichment of J2 in FUS-P525L-GFP granules (I) and in double positive granules (K). (G-K) Mann-Whitney test, n=3.

Mitochondria are known to be a source of dsRNA production in the cell, as sense and antisense transcripts of the circular genome can align to form dsRNA, which can trigger an MDA5-dependent lethal IFN1 response when released into the cytosol ^21^. We therefore measured dsRNA abundance using J2 immunofluorescence staining. We found a clear J2 signal inside and outside the mitochondria in FUS^wt^ and FUS^mut^ sMNs (Fig. 2B-D Supplementary Fig. 1B). At baseline, we observed a significantly higher amount of mitochondrial J2 in FUS-P525L mutant sMNs. Notably, we also identified a higher proportion of cytosolic J2-dsRNA outside the mitochondria in FUS^mut^ sMNs (Fig. 2C-D). Another important finding was the translocation of J2 to FUS granules. Consistent with previous reports, FUS-ALS sMNs spontaneously produced FUS-GFP granules (Fig. 2F). Double immunostaining revealed that ∼45% of FUS-GFP granules were double labelled with the stress granule marker G3BP1 (Fig. 2F). While there was no difference in the proportion of G3BP1+/FUS-gfp+ granules (Fig. 2H), FUS P525L sMNs showed significantly more FUS-gfp+ granules (Fig. 2G). Notably, FUS P525L sMNs contained more dsRNA-containing granules, both within G3BP1+/FUS-gfp+ granules and within G3BP1-/FUS-gfp+ granules (Fig. 2 I-K). Subtraction of this granule J2 signal from extramitochondrial J2 yielded the free cytosolic J2 fraction (Fig. 2E), which was significantly higher in FUS^mut^ sMNs. In conclusion, we show that FUS^mut^ sMNs have significantly increased dsRNA in mitochondria, SGs, but also released into the cytosol of untreated FUS^mut^ sMNs.

### ISG activation is mediated by RIG-I signaling

Since we found significantly increased dsRNA in FUS^mut^ sMNs, we sought to further investigate the role of the cytosolic dsRNA sensors MDA5 and RIG-I in FUS^mut^ sMNs. Remarkably, we did not find a detectable signal for MDA5 on WB in either FUS^wt^ or FUS^mut^ sMNs. Since both sensors are ISGs themselves, stimulation of the pathway with IFN-β was used as a positive control. A clear band was observed at the expected 140kDA (Supplementary Fig. S1C). The same was true for RIG-I, which showed a strong signal at 100 kDA after exposure to IFN-β. Notably, however, a distinct band for RIG-I was already detectable in FUS^mut^ sMNs under control conditions (Fig. 3C,D; Supplementary Fig. S2C). Consistent with this, we also observed increased expression of RIG-I1 in FUS^mut^ sMNs in immunofluorescence staining (Fig. 3A) and significantly increased expression by qPCR (Fig. 3E). Notably, RIG-I was also highly expressed in the axons of motor neurons (Fig. 3B).

**Figure 3.**
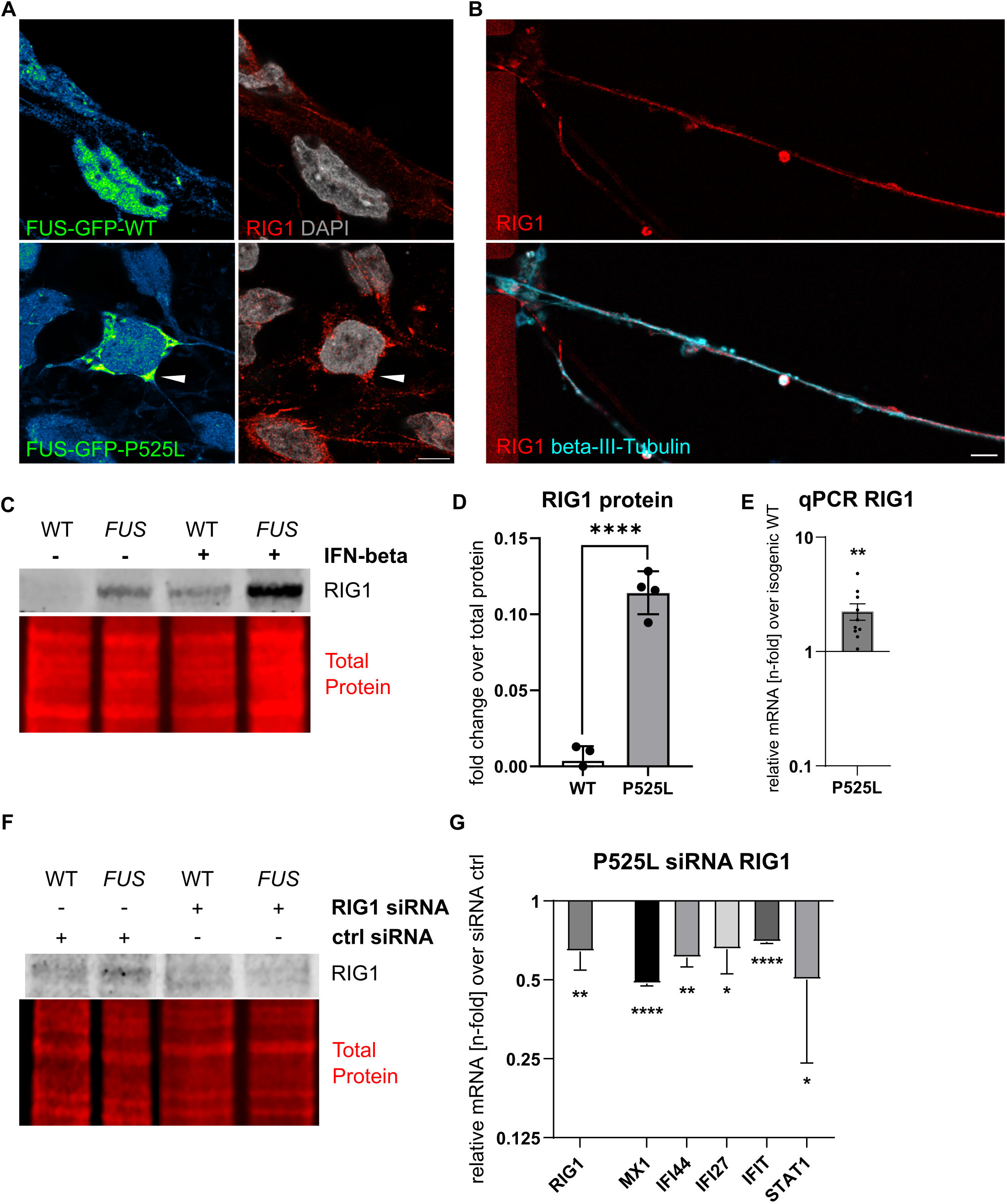
Expression of RIG-I in FUS-P525L sMNs. (A, B) IF panel of FUS^wt^ and FUS^mut^ sMNs counterstained for RIG-I. Scale bar 10µM. Note the axonal localization and enrichment in FUS-GFP granules (white arrowhead). (C) Western blot scan of RIG-I (100kDA) in FUS^wt^ and FUS-P525L sMNs. IFN-beta treatment (20ng/ml 24h) was performed as a positive control. (D) Quantification of (C) normalized to a total protein staining, n=4, unpaired t-test. (E) Significantly higher RIG-I mRNA expression was found in P525L-sMN compared to control by RT-qPCR, n=10, unpaired t-test. (F) WB scan of RIG-I for WT and FUS-P525L sMNs treated with either ctrl-siRNA or RIG-I siRNA for 3d. Note the reduced signal of RIG-I in FUS^mut^ sMNs following RIG-I siRNA. (G) RT-qPCR in sMNs FUS-P525 samples treated with siRNA against RIG-I for 3d normalized to ctrl siRNA treated samples. Modest reduction of depicted ISGs including RIG-I, unpaired t-test, n=3.

We next asked whether RIG-I activation was a cause of TBK1 stimulation and ISG activation or a consequence of being an ISG itself. To address this, we performed siRNA-mediated knockdown of RIG-I. Liposomal-based siRNA treatment over 72h resulted in a reduction of RIG-I at the protein level (Fig. 3F) and a robust decrease of RIG-I mRNA in sMNs (Fig. 3G). This was accompanied by a marked decrease in mRNA for the ISGs MX1, IFI27 and IFI44 (Fig. 3G). Taken together, these data suggest that ISG activation in the FUS mutation is mediated by RIG-I activation.

### RIG-I is increased in postmortem spinal motor neurons of FUS patients

To further validate these in vitro findings, we analyzed RIG-I expression in spinal cord FFPE tissue section of FUS-ALS patients compared to healthy controls. While there was a faint cytoplasmic staining of RIG-I-I in healthy control α-MNs, there was a marked elevated staining in case of FUS-ALS patients (Fig. 4).

**Figure 4.**
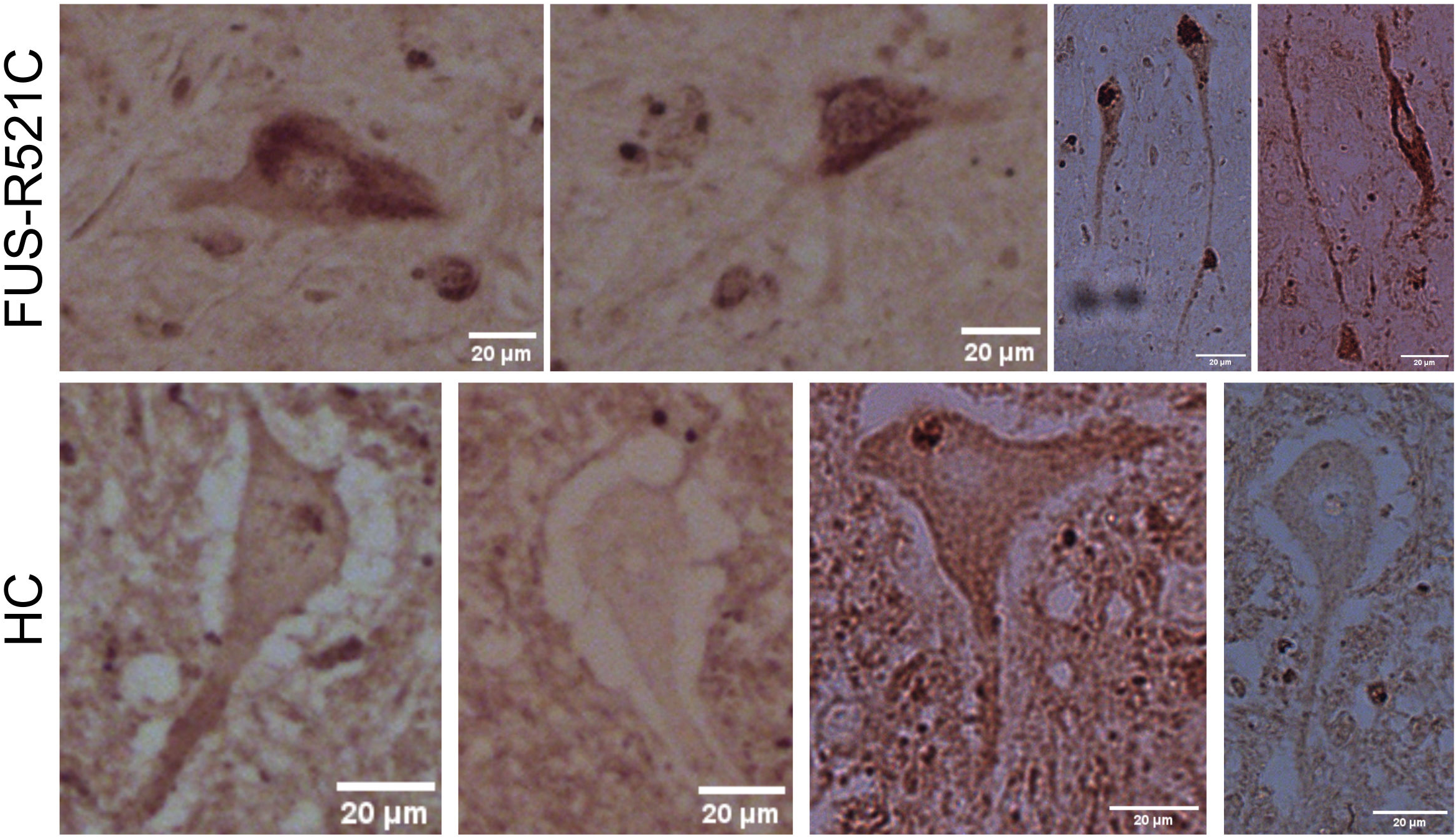
IHC for RIG-I of spinal cord sections of a FUS-ALS patient with a FUS-R521C mutation and a healthy individual. Note the higher staining intensity in sMNs of the FUS-ALS patient, which was partially particularly intensive in axons whereas the sMNs in the healthy individual showed rarely small inclusions.

### Interferon signaling induces axonal degeneration

Considerable evidence has accumulated that DNA damage might be upstream of (axonal) neurodegeneration in different ALS models, including FUS^13,16^. However, the link between the two remains to be elucidated. Previous reports have demonstrated functional axonal deterioration and neurodegenerative phenotypes after exposure of neurons to TNF-α or IFN-β ^17,18^. In light of these previous findings, we asked whether IFN1 activation might be responsible for axonal damage and neurodegeneration. Therefore, we exposed mature motor neurons to IFN-β for 7 days and assessed the axonal growth area by brightfield imaging before and after treatment (Fig. 5 A-D). For selective axonal analysis, we used microfluidic chambers, which allow the cultivation of neuronal somata spatially separated from their outgrowing axons, as previously shown ^13^. While WT axons showed marked growth over the seven days as measured by the area ratio normalised to the area just before treatment, the growth of FUS^mut^ axons was significantly reduced (Fig 5 A,B). Treatment with IFN-β 20ng/ml reduced axonal area in WT neurons but had no additional effect on FUS^mut^ sMNs axons (Fig. 5C). To validate this structural phenotype, we measured neurofilament light chain (NfL) in the media supernatant as a surrogate for axonal damage and a well-established diagnostic and prognostic biomarker in the clinic ^22^. Medium of mature sMNs was collected after three days of IFN-β or mock treatment and measured using ELLA. First, we observed a strikingly higher NfL amount in FUS^mut^ vs. FUS^wt^ sMNs under baseline conditions (Fig. 5F). Importantly, IFN-β led to a significant NfL increase in FUS^wt^ sMNs, while mutant sMNs did not change. These results suggest that IFN signaling is harmful for sMNs leading to neurodegeneration and even more, that IFN1 signaling cascade might be one mediator of the observed axon degeneration in FUS-ALS sMNs.

**Figure 5.**
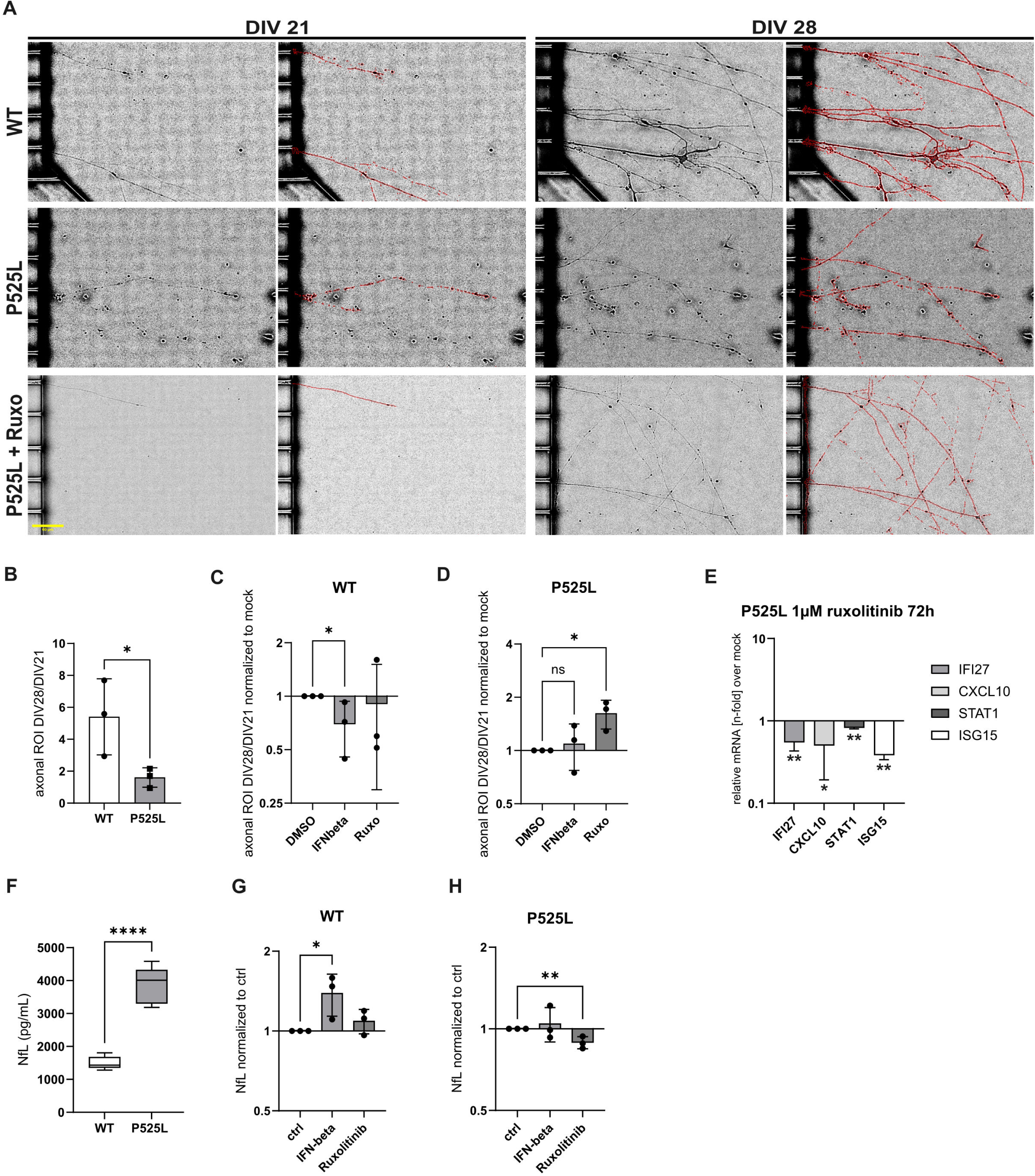
JAK Inhibition can partially reverse phenotypes in FUS sMNs. (A) Brightfield image panel (20x) showing the axonal compartment of the microfluidic chambers in which sMNs were cultured on the left (proximal) side. Subsequent imaging was performed after 7 days and image segmentation to detect axons was performed with Fiji as indicated by the red ROI area. Scale bar 50µM, yellow, bottom left. (B) Quantification of the ROI area ratio (ROI DIV28/DIV21) indicating a significantly lower axonal growth in FUS-P525L sMNs compared to control. N=3, unpaired t-test. (C, D) Axonal ROI ratio in FUS^WT^ (C) or FUS-P525L (D) sMNs treated with IFN-beta 20ng or ruxolitinib 1µM for 7d normalized to respective mock control, unpaired t-test, n=3, unpaired t-test. (E) RT-qPCR for a set of ISGs in FUS-P525L sMNs treated with ruxolitinib for 72h. Normalized to DMSO-treated controls there was a partial reduction of the depicted ISGs, unpaired t-test, n=3. (F, G, H) Neurofilament light chain (NfL) was measured in medium supernatant by ELISA. Medium was collected after 72h on DIV21 from FUS^wt^ and FUS-P525L sMNs. Cells were treated with either IFN-beta 20ng/ml or ruxolitinib 1µM for 3d and NfL levels were normalized to the respective control condition (DMSO for ruxolitinib and PBS for IFN-beta). N=3, unpaired t-test.

### Axonal damage can be mitigated by JAK Inhibition

IFN1 signaling activates the JAK-STAT pathway, for which FDA-approved inhibitors are available. Therefore, we exposed mature sMNs to the JAK inhibitor ruxolitinib for 7 days and assessed the axonal growth area by brightfield imaging before and after treatment (Fig. 5 A-D). While WT axons showed no change in their outgrowth behavior in the presence of ruxolitinib, FUS^mut^ axonal growth could be significantly restored by treatment with 1 µM ruxolitinib for seven days (Fig. 5 D). Ruxolitinib also significantly reduced ISGs in FUS^mut^ sMNs (Fig. 5E). Medium from mature sMNs was collected after three days of ruxolitinib or sham treatment and measured using ELLA. Ruxolitinib resulted in a highly significant reduction of NfL in FUS^mut^ sMNs, but no changes in FUS^wt^ sMNs (Fig. 5G,H). These results support the idea of an existing activation of the IFN pathway in FUS^mut^ sMNs, but not in FUS^wt^ sMNs, which can be alleviated by JAK inhibition.

### Upregulated mitochondrial transcription in axonal FUS-P525-sMN samples

Having shown increased cell-autonomous IFN1 signalling in sMNs of FUS-ALS patients responsible for axonal degeneration together with prominent axonal RIG-I expression (Figure 3), we wondered whether the source of increased extramitochondrial dsRNA was indeed increased mitochondrial transcription. To this end, we used our previously described RNA sequencing (RNA-seq) dataset in which we performed spatial transcriptomics of somatodendritic vs. axonal compartments of FUS^wt^ vs. FUS^mut^ sMNs using microfluidic devices ^23^. To identify possible protein-protein interaction networks, we performed STRING analysis on the axon-specific differentially expressed genes (DEGs) between FUS^wt^ and FUS^mut^ datasets. Notably, using this unbiased sequencing approach, we identified an axonally upregulated gene cluster involved in RIG-I-based innate immune activation (Figure 6B). Furthermore, we found a protein-protein interaction network of axonally enriched genes of mitochondrial/metabolic functions (Fig. 6A). Mitochondrial transcription is carried out by the single subunit mitochondrial RNA polymerase (POLRMT). We therefore wanted to investigate whether increased mitochondrial transcription could be the source of ISG activation. We used the non-competitive human mitochondrial RNA polymerase (POLRMT) inhibitor IMT1 ^24^. IMT1 inhibition was shown previously to lower dsRNA in human cells ^25^. This led to a dose-dependent drop of ISGs in FUS-P525L sMNs (Fig. 6C). These data suggest that an increased mitochondrial transcription in FUS^mut^sMNs led to excessive dsRNA production, which triggers the innate immune system via RIG-I.

**Figure 6.**
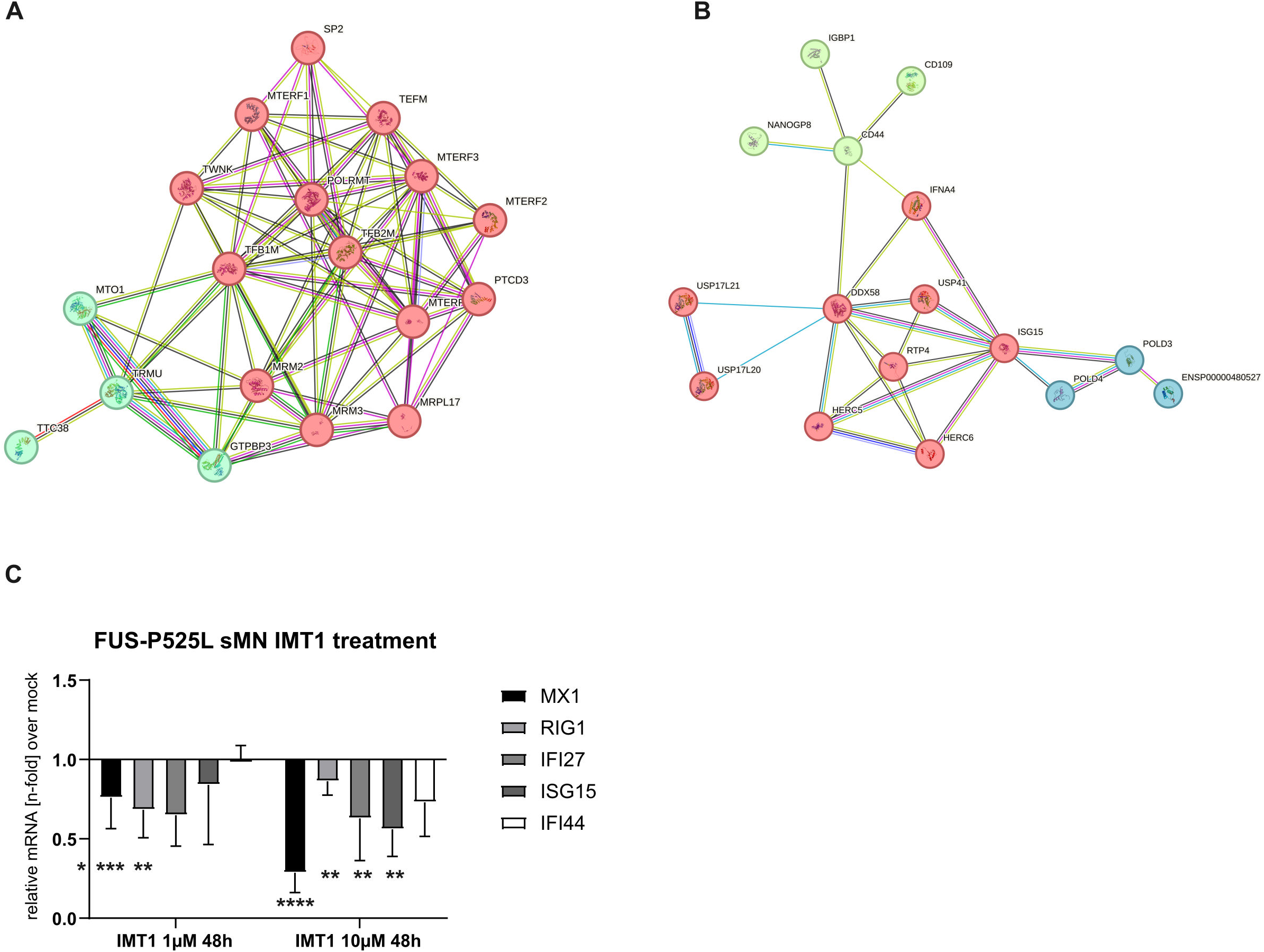
Axonal RNA sequencing reveals upregulated mitochondrial transcription in axonal FUS-P525 sMNs samples. Protein-protein interaction based on transcriptome analysis from RNA-seq data of axonal sMNs samples. (A) Cluster indicates significant upregulation of innate immune activation with RIG-I in the center and (B) mitochondrial transcription pathway enrichment. (C) Dose-dependent drop of ISGs in FUS-P525L sMNs after treatment with the non-competitive, human mitochondrial RNA polymerase (POLRMT) inhibitor IMT1, One-Way ANOVA, Tukey’s post hoc, n=3.

### IFN1 activation is found in peripheral blood samples of FUS-ALS patients

Our previous results showed a significant neuron-specific upregulation of IFN1 signalling in various patient-derived spinal motor neurons and post-mortem spinal cord from FUS-ALS patients. However, we also wondered whether these findings might be of clinical relevance. We therefore analysed ISG upregulation using the Interferon Signature Score (ISS), which combines a number of ISG measured by qPCR in peripheral blood samples (Fig. 7A,B). This assessment of IFN1 upregulation is widely used in rheumatology and correlates with disease activity in inflammatory diseases such as systemic lupus erythematosus ^26^.

**Figure 7.**
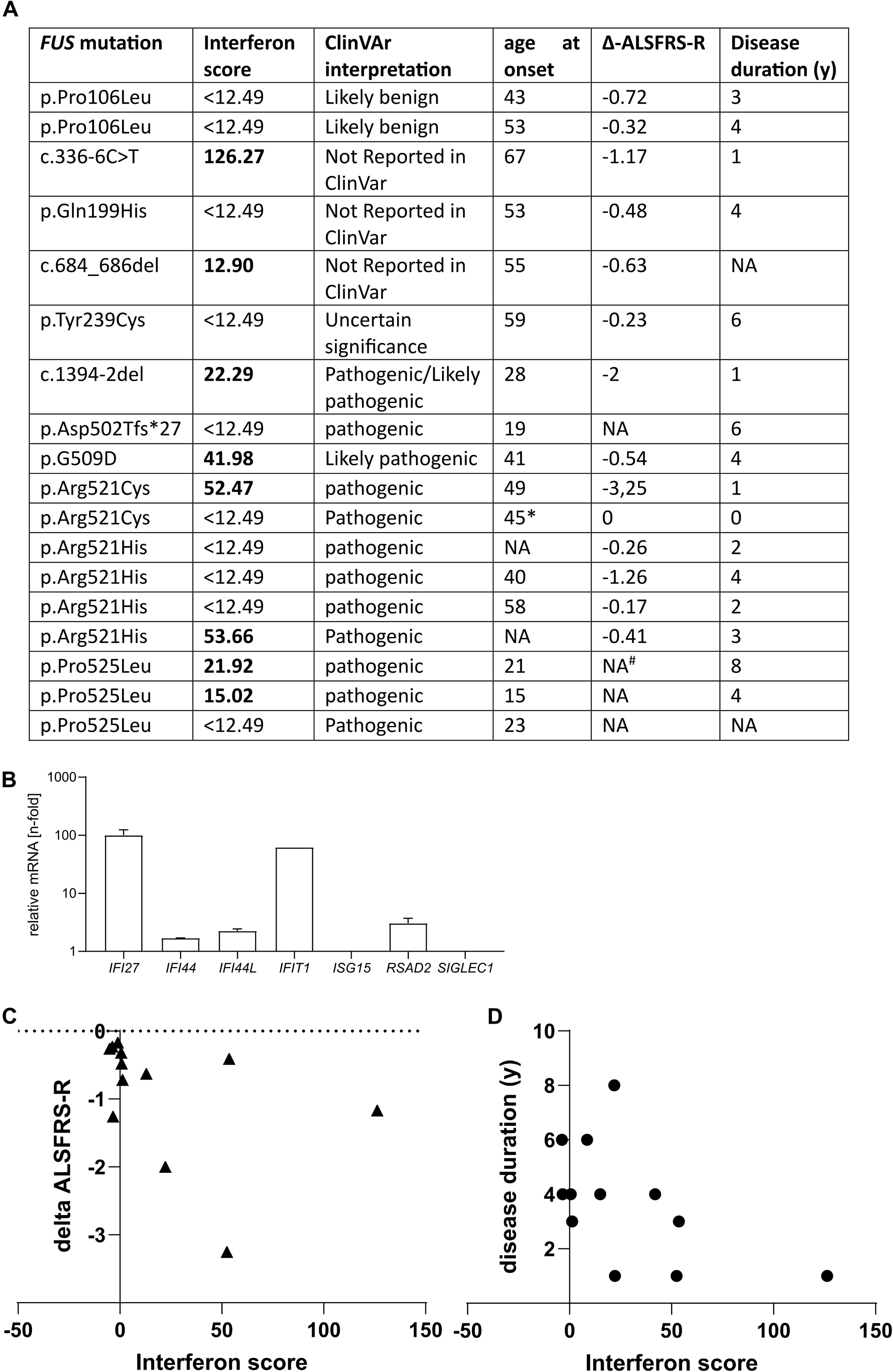
IFN score assessment in blood samples of FUS-ALS patients. (A) Analysis of the interferon stimulated gene response in 18 ALS patients with mutations in FUS including assessment of clinical parameters. Seven mRNA transcripts were analyzed as indicated in (B) with qPCR normalized to HPRT and GAPDH. The interferon signature was calculated as described before ^50^. An IFN score of 12.49 indicates the cut-off for significant results and is calculated as the median of 10 healthy controls plus 2.5xSD. Any IFN score higher than 12.49 is considered increased. (B) Example of one blood ISG transcription measurement in a FUS-ALS patient with a FUS c.336-6C>T mutation, which has not been reported previously. (C) A Significant correlation was found between the Δ-ALSFRS-R and the IFN score (Spearman r = -0,5879, p =0.038, one-tailed t-test). * pre-symptomatic patient with pure fasciculations and signs of denervation on EMG. ^#^ patient was locked in for several years at the time of the blood drawl.

We thus investigated 18 FUS-ALS patients with different FUS mutations for an interferon signature (Fig. 7). We performed RT-qPCR to calculate the interferon score, which is considered to be increased if greater than 12.49, which is the median + two-fold SD of a healthy control group. Interestingly, eight out of 18 FUS-ALS patients showed an increased interferon score (Fig. 7A). Notably, only two patients with an N-terminal mutation had an elevated interferon score, whereas 50% of C-terminal mutation carriers had an elevated score. We further correlated the interferon score with several clinical variables. We found a significant correlation of the ISS with the delta-ALSFRS-R, which is a measure of disease progression (Spearman r= -0.59, p=0.024, Fig. 7C) and disease duration (Spearman r=-0.58, p=0.04), but not with age at onset (Spearman r=-0.49, p=0.08).

## Discussion

The vast majority of clinical trials in ALS fail by phase 3. One reason may be the heterogeneous nature of ALS and the lack of pathophysiological-driven biomarkers for patient stratification. IFN1 pathway activation, an emerging field in ALS research, may provide an opportunity to fill this gap. It is also of interest because it culminates in a pathway for which there are already FDA and EMA approved drugs, irrespective of the different upstream mechanisms involved in its activation. Accordingly, IFN1 activation has been reported in various genetic ALS model systems and patients, mainly involving cGAS STING-dependent pathways, although it appears that in ALS there is a plethora of upstream mechanisms to trigger these pathways depending on specific mutations ^4,10–12,27^.

Since the pathophysiology of FUS-ALS includes impairment of DNA damage repair as well as mitochondrial damage, we hypothesised that one of these processes might be involved in aberrant IFN1 activation. Using isogenic and non-isogenic iPSC-derived sMNs as well as post-mortem spinal cord tissue, we were able to demonstrate an IFN1 pathway activation in FUS-ALS, which was dependent on RIG-I activation. RIG-I was increased in sMNs axons of FUS^mut^ iPSCs as well as in α-MNs of FUS-ALS post-mortem spinal cord, presumably through increased mitochondrial transcription. Notably, IFN signalling induced axonal degeneration in sMNs. Finally, ISG activation was found in the peripheral blood of half of FUS-ALS patients and correlated with disease aggressiveness and duration. Janus kinase inhibition by FDA-approved ruxolitinib reversed not only ISG signaling but also axonal damage and neurofilament release of diseased neurons, supporting it as a candidate as a biomarker-driven target for (FUS-)ALS.

While most studies of IFN1 activation have been conducted in non-neuronal cells, including immune cells, only a few recent reports have implicated neuronal STING activation in ALS and FTD, including analysis of post-mortem cortical and spinal cord ^2,4,12^. The in vivo data remarkably showed that this STING activation was most prominent in the most affected neuronal subpopulations in ALS, namely cortical layer V and spinal α-sMNs ^12^. Furthermore, pathway components were found intra-neuronally, suggesting the cell-autonomous nature of activation with DNA damage as the main contributor. However, FUS-ALS was not included in this study. Interestingly, our results show that cell-autonomous IFN1 activation is also present in FUS-ALS sMNs, although not primarily induced by nuclear DNA damage and subsequent STING activation. This was supported by the lack of IFN1 activation after DNA damage induction and the lack of efficacy of the STING inhibitor H151 in reducing ISGs in the FUS^mut^ sMNs.

In contrast, we were able to delineate the importance of RIG-I activation in FUS^mut^ sMNs to trigger the innate immune response. Consistent with the role of RIG-I in sensing dsRNA species, we found abundant dsRNA in the cytosol of FUS^mut^ sMNs, where RIG-I resides. Consistent with this, our RNA-seq data indicated an up-regulation of mitochondrial RNA transcription pathways that could promote the accumulation of immunogenic dsRNA in FUS-P525L sMNs. Notably, pharmacological inhibition of mitochondrial transcription with IMT1 ^24^ could reduce ISG expression in FUS-P525L sMNs, further supporting the mechanism of dsRNA-mediated innate immune stimulation in our FUS sMNs model. However, future studies are needed to elucidate the source and fate of immunogenic dsRNA in sMNs. One could speculate on mechanisms that prevent aberrant innate immune activation by dsRNA in neurons. One such mechanism could be the integration of dsRNA into stress granules, the absence of which has been reported to lead to hyperactivation of IFN type 1 ^28,29^. However, the binding of viral RNA and dsRNA to G3BP1 has been reported to trigger an IFN1 response mediated by RIG-I ^30^. Interestingly, SG assembly was reported to be required for IRF3-mediated IFN production, but not IFN signaling or proinflammatory cytokine induction ^31^. In FUS-ALS, cytoplasmic mislocalization is reported to induce SGs and these show aberrant dynamics ^32^, thus SG function might be disturbed, which might indeed trigger IFN1 response activation. Furthermore, compensatory mechanisms such as heat shock factor expression or integrated stress response activation are already increased under baseline conditions ^32^, thus any additional event affecting SGs might be sufficient to neutralize SG’s protective effect. Further work is however needed to clarify the role of SGs in dsRNA-RIG-I mediated IFN1 activation and particularly in FUS-ALS.

Given reports of both physiological functions for FUS in mitochondrial DNA repair ^33^ and pathological accumulation of mutant FUS leading to HSP60 dysfunction^34^, it will be important to investigate the role of FUS with a focus on mitochondrial dsRNA maintenance, as both loss-of-function and gain-of-function mechanisms could apply. Mechanistically, our experiments support a RIG-I-driven sensing of the abundant cytosolic dsRNA in FUS^mut^ sMNs and subsequent innate immune stimulation. In contrast, we did not find a meaningful signal for MDA5, another established dsRNA sensing enzyme upstream of TBK1 activation. Our data also revealed enrichment of RIG-II in post-mortem sMNs from FUS patients, further supporting the critical role of this pathway in our experiments. Importantly, accumulation of endogenous RNAs as a result of TDP-43 loss of function has previously been shown to cause RIG-I-dependent necrotic cell death (Dunker, Ye et al. 2021). Post-mortem evidence for neuronal cell-autonomous innate immune activation already exists for both, sporadic and genetic ALS models ^35, 12, 36^, but highlighted the role of STING signaling in this. This has been experimentally attributed to upstream DNA damage events leading to STING activation in vitro. However, this does not contradict our data, as STING itself has been shown to be an ISG ^37^. Therefore, the strong post-mortem signal in sMN in the presented singular FUS-ALS case ^12^ could be due to primary RIG-I signaling and secondary STING stimulation as an ISG, which could also support our finding of slightly, but not significantly increased STING on the WB. Consistent with this, the STING inhibitor H151 failed to reduce ISGs in FUS^mut^ sMNs in our experiments. This would support the difference between a possible upregulation by innate immune stimulation and the primary activation by cGAS, which should be blocked by H151.

Regardless of the primary mechanism, the common downstream canonical TBK1-IRF3 signaling pathway is a promising drug target. Our data and others ^9^ suggest that drugs such as ruxolitinib, whose JAK inhibitory mode of action covers a broad spectrum of IFN1 signaling effects, could not only reduce ISG and NfL levels in FUS^mut^ sMNs but also improve axonal outgrowth in our cell model. The non-canonical NF-κB pathway, which is thought to be activated secondary to TBK1 phosphorylation of p65, may play an important role in a neurodegenerative phenotype by stimulating the expression of other pro-inflammatory factors such as TNFα ^38^. As we also observed p65 phosphorylation in FUS-P525L sMNs at the Ser-536, the target site for TBK1 ^39^, this represents another promising mechanism to evaluate for pharmacological intervention in future studies. Importantly, although there is evidence for both, IRF3 and NF-κB signaling following STING ^40^ or RIG-I/MAVS ^41–43^ activation, further studies are required to elucidate the dominant pathophysiological mechanisms occurring in (spinal moto)neurons.

While both, impairment of DNA damage repair and axonal mitochondrial trafficking deficits and depolarization have been reported as central phenotypes in FUS-ALS, their connection so far remained enigmatic ^13,15,44^. While DNA damage induction was sufficient to phenocopy FUS-ALS mitochondrial deficits ^13^, it did not induce an IFN1 response. However, IFN-β treatment was indeed able to induce axon degeneration (Fig. 5). Mutant FUS was reported to localize to mitochondria, being involved in mitochondrial DNA damage repair ^33^. Since we and others recently showed that increased DNA damage leads to activation of DNA-PK and this results in further increase of cytoplasmic mislocalization of FUS putatively due to its phosphorylation closing a vicious cycle ^13,45^, we speculate that the increased cytoplasmic FUS increasingly culminates in mitochondria and stress granules, both of which are involved in RIG-I mediated IFN1 activation. The latter then becomes suicidal, including the observed axon degeneration. However, this hypothesis requires further investigation.

Another limitation of the study is that no CSF was available to analyze the IFN1 response in the CSF of FUS-ALS patients. Of note, however, ISG activation by the qPCR would be difficult to assess in CSF due to the low cell numbers in non-inflammatory neurodegenerative diseases. Alternatively, ISGs such as CCL2 have been reported to be increased at the protein level in ALS CSF, which was not always seen in serum, so sensitivity may be even higher when analysing CSF ^46^. In addition, the source of RIG-I activation requires further investigation. In addition, the role of increased cytosolic DNA and the cGAS-STING pathway and its putative crosstalk with the RIG-I pathway in FUS-ALS requires further investigation. Finally, future studies are needed to systematically and differentially assess body compartment differences in IFN1 activation in ALS and correlations with neuropathology-driven biomarkers such as cryptic peptides ^47^ or extracellular vesicle TDP43 ^48^.

## Methods

### Cell Culture

iPSC-derived sMNs culture was accomplished as demonstrated previously ^13,32^. The iPSC generation and CRISPR-Cas9n creation of the isogenic pair of lines of hiPSC and NPC of WT FUS-eGFP and P525L FUS-eGFP is described therein including the generation of the other cell lines carrying FUS mutations (R521C, R521L, R495Qfs*527) and two healthy, non-isogenic control lines. Note, that the non-isogenic FUS^mut^ cell lines were only used for experiments demonstrated in Figure 1. NPC culture and differentiation was performed via a modified protocol by Reinhardt et al. ^49^. Briefly, NPC were maintained in the basic medium (DMEM-F12/Neurobasal 50:50 medium with N2 Supplement (1:200), B27 Supplement without vitamin A (1:100), penicillin/streptomycin/glutamine (1%), supplemented with Chiron-99021 (3 μM), Ascorbic acid (150 μM) and Purmorphamine (0.5 μM) on dishes coated with Matrigel. For coating, Matrigel was diluted 1:100 in Knock-Out DMEM and added to dishes, followed by the incubation at 37°C for at least 1 h. NPC were split after reaching 70–80% confluence at a 1:10 ratio using Accutase for 10 min at 37°C. Induction of sMNs differentiation was performed by adding basic medium supplemented with BDNF (1 ng/mL), Ascorbic acid (200 μM), Retinoic acid (1 μM), GDNF (1 ng/mL) and Purmorphamine (0.5 μM) to freshly split NPCs and maintained for 5 days with the medium changed every other day. For the final maturation, the medium was changed on day 6 to the basic medium supplemented with DBcAMP (100 μM), BDNF (2 ng/mL), Ascorbic acid (200 μM), TGFβ-3 (1 ng/mL) and GDNF (2 ng/mL). On day 9, cells were split onto the dishes coated with Poly-L-ornithine and laminin and maintained for at least 21 days (DIV21) before they were used for the final analysis. In case of the brightfield experiments shown in Figure 5, cells were kept for an additional 7 days in culture (DIV28). Furthermore, in this experiment cells were grown in microfluidic chambers (MFC) allowing selective axonal assessments. MFC culture for iPSC-derived sMNs was demonstrated previously by us ^13^. The cells were regularly tested for mycoplasma contamination.

### Patient characteristics

Cell lines carrying a clinically more benign phenotype (R521C, L) or more severe (R495QfsX527, P525L) FUS mutation were included in this study and systematically compared to two control iPSC lines from healthy volunteers or to an isogenic line. All procedures were in accordance with the Helsinki Convention and approved by the Ethical Committee of the University Medical Center Rostock (A2019-0134) and Dresden (EK45022009) Patients and controls gave written consent before skin biopsy or blood sampling for ISG measurement. The delta-ALSRFS-R was calculated as the ratio of the maximum ALSFRS-R (48) minus the ALSFRS-R at the point of blood drawl divided through the estimated amount of months since occurrence of first ALS symptoms.

### Treatments and siRNA

Etoposide (Sigma-Aldrich E1383) was dissolved in DMSO to obtain a 10 mM stock. ABT-888 (PARP1 inhibitor, Santa Cruz Biotechnology sc-202901) was dissolved in DMSO to obtain a 20 mg/ml stock. Recombinant human interferon-beta (PeproTech, 300-02BC) and recombinant human TNF-alpha (PeprotTech, 300-01A) were resuspended to a final stock of 20µg/ml in sterile aqua dest. Ruxolitinib was obtained from MedChemExpress (HY-50856) and dissolved in DMSO for a 10mM stock. The STING inhibitor H151 was obtained from Tocris (6675) and dissolved in DMSO to a stock of 5 mM. Thapsigargin was from Sigma (T9033), dissolved in DMSO, and kept at -80°C. IMT1 was purchased from MedChemExpress (HY-134539) and dissolved in DMSO. siRNA Silencer Select was from Invitrogen (DDX58/RIG-I 4392420; Negative Control 4390843). Silencer Select siRNA was dissolved in nuclease-free water according to the manufacturer’s instructions. Transfection was done using FUSE-it siRNA (beniag GmbH) according to the manufacturer’s instructions. In brief, the master mix was produced according to the manufacturer’s instructions. The sMNs were then incubated for 20 min in an incubator at 37°. RNA or protein isolation was performed after 72h.

### Microscopy and Immunofluorescence

Cells were fixed in vitro with ice-cold 4% paraformaldehyde for 151minutes, then permeabilized in 0.2% Triton X-100 for 101minutes. Afterward, cells were washed three times with PBS for 51minutes and blocked in Pierce Protein-free blocking buffer (37572; Thermo Fisher) for 1 hour at room temperature. The following primary antibodies were diluted in Pierce Protein-free blocking buffer and incubated overnight at 4°C: Anti-RIG-I (1:500, clone 1C3, MABF297 Sigma-Aldrich), anti-dsRNA J2 (1:500, SCICONS 10010200); IgM-anti-DNA 30-10-10 (1:500, Progen 690014), anti-TOM20 (1:1000, Invitrogen, MA5-34964), anti-HSP60 (1:1000; ab46798; Abcam). Afterward, cells were washed thrice in PBS for 51minutes, secondary antibodies were diluted 1:1000 in blocking buffer and incubated for 11hour at room temperature, on a shaker. Following three more washing steps with PBS for 5 min, cells were mounted in DAPI Fluoromount-G mounting medium (0100-20; Southern Biotechnology). Images were acquired on a Zeiss inverted AxioObserver.Z1 microscope with LSM 900 module and high-resolution Airyscan 2 module, using a 63×11.4 NA plan apochromat objective.

### Image Analysis with Fiji

CZI images were imported and edited in FIJI (2.15.1). The Labkit plugin (15) was applied for image segmentation to generate defined classifiers for, e.g., the fluorescence signal of dsRNA. From this, ROIs were created for each signal of interest. Using logical operators the nuclear area was removed and the area proportion of dsRNA inside and outside of the HSP60+ ROI was quantified. This was normalized to the total cell area measured with a background GFP classifier.

For MFC brightfield quantification, the total area of the right MFC compartment was captured with a 20x objective and repeated imaging with an image overlay of 10%. Image fusion was accomplished with a Zeiss ZEN Blue built-in processing algorithm. Axonal segmentation for ROI recognition was performed with the Labkit plugin to capture the entire axonal growth area within the dish.

### Statistics

Statistical testing was performed in GraphPad Prism 10. Statistical tests are described in the figure captions. Asterisks indicate significance in figures as a result of statistical testing: * p<0.05, ** p<0.01, ***p< 0.001, **** p<0.0001.

### Western blot

Cell pellets were scraped from the dish and snap-frozen. Afterward, they were lysed in RIPA buffer containing EDTA-free protease inhibitor cocktail (04693132001; Roche) and phosphatase-inhibitor (Roche PhosSTOP EASYpack 04906837001) for 301minutes on ice. The lysate was incubated with Roti-Load (Carl Roth, K929.1) and heated for 5 minutes at 96°C. A total of 30 μg of protein was loaded on precast polyacrylamide gels (Bio-Rad 4-15% Criterion TGX Stain-Free Precast Gels, 18 Well Comb, 30 µl, 1.0mm, 5678084) for gel electrophoresis. Proteins were blotted on 0.2-μm nitrocellulose membrane (Trans-Blot Turbo Midi 0.2 µm Nitrocellulose Transfer Packs 1704159; BioRad), using a Trans-blot Turbo transfer system (221V, 71minutes; Bio-Rad). Total protein was stained with REVERT Total Protein Stain (Li-Cor: 92611011) according to the manufacturer’s instructions. Membrane blocking was done in 5% non-fat milk in TBS-T (T145.3; in TBS; Roth) for 1 hour at room temperature. Primary antibodies were diluted in blocking solution and kept on the membrane for 1 hour at room temperature if not stated otherwise. For WB the following primary antibodies were used: rabbit anti-MDA-5 (Cell Signaling, D74E4); rabbit anti-TBK1 (Invitrogen, PA5-17478), rabbit anti-RIG-I (D14G6,Cell Signaling, #3743), rabbit anti-phospho-TBK1 (Ser172, D52C2, Cell Signaling, #5483), mouse anti-IRF3 (Invitrogen, 14-9947-82 eBioscience), rabbit anti-STING (Novus, NBP2-24683), rabbit anti-p65 (C22B4, Cell Signaling, #4764), rabbit anti-phospho-Ser536-p65 (93H1, Cell Signaling, #3033). Secondary antibodies (anti-rabbit IRDye 800CW abcam Ab216773, anti-mouse IRDye 800CW abcam ab216772, anti-rabbit IRDye® 680RD abcam ab216777, anti-mouse IRDye® 680RD abcam ab216776) were incubated in TBS-T overnight at 4°C. Images were acquired on a Li-COR ODYSSEY XF analyzer. Image analysis was performed with the Empiria Studio 3.0 software (Li-COR).

### Quantitative RT-PCR

RNA was extracted from isolation cell pellets and isolated with the Quick-RNA Miniprep Plus Kit (Zymo Research R1057T) according to the manufacturer’s instructions. cDNA was generated with the High-Capacity cDNA Reverse Transcription Kit (ThermoFisher 4368814) using 200ng of isolated RNA according to the manufacturer’s instructions. Gene expression was determined by quantitative rt-PCR using the FastStart Essential DNA Green Master kit (06402712001) in a LightCycler 480 II (Roche) and normalized to the mean of the 18s and HPRT expression. The following primer sequences were used in this work: 18s fwd CGTAGTTCCGACCATAAACGATGCC and rev GTGGTGCCCTTCCGTCAATTCC, HPRT fwd CCTCCTCCTCTGCTCCGCCA and rev GGTTCATCATCACTAATCACGACGCCAG, CXCL10 fwd CCTGCATCAGCATTAGTAATCAACC and rev TGGATTCAGACATCTCTTCTCACC, IFIT1 fwd CCTTGCTGAAGTGTGGAGGA and rev CCTGCCTTAGGGGAAGCAAA, IFI27 fwd TGCTACAGTTGTGATTGGAGG and rev ACTGCAGAGTAGCCACAAGG, MX1 fwd CCAGCTCAGGGGCTTTGG and rev TTGGAATGGTGGCTGGATGG, STAT1 fwd AGTGTAAGTGAACACAGAAGAGTC and rev GTAACACGGGGATCTCAACAAG, RIG-I fwd TGAAGCCATTGAAAGTTGGG and rev CCATCATCCCCTTAGTAGAGC, SIGLEC1 fwd ACCTGGAGGAAACTGACAGTGG and rev CTCAGTGTCACTGCCTGTCCTT, RSAD2 fwd CCAGTGCAACTACAAATGCGGC and rev CGGTCTTGAAGAAATGGCTCTCC, ISG15 fwd CTCTGAGCATCCTGGTGAGGAA and rev AAGGTCAGCCAGAACAGGTCGT, IFI44L fwd and rev, IFI44 fwd TCTATTCAATACTTCTCCTCTCAGATGATAG and rev TGAGCAAAGCCACATGTACCA.

The interferon signature score was calculated as demonstrated previously ^50^. An IFN score above the 2.5x SD of the control group (> 12.49) was considered increased.

### Neurofilament in supernatant (ELLA)

NfL levels in iPSC supernatants were measured with the Ella microfluidic system using the Simple Plex Human NF-L Cartridge (BioTechne, Minneapolis, USA) according to the manufactureŕs instructions. Quality control samples were included in all runs to monitor assay performance.

### Post mortem

Postmortem human tissue was obtained from ALS patients at the department of Neuropathology of the Amsterdam UMC (University of Amsterdam, the Netherlands). All patients with ALS fulfilled El Escorial criteria for diagnosis, as reviewed independently by two neuropathologists. ALS subjects died from respiratory failure or euthanasia. Tissues obtained from patients, who died because of non-neurological diseases were used as controls. No signs of infection before death were detected in both ALS and control patients included in the study. Informed consent was obtained for the use of brain tissue and access to medical records for research purposes; approval was obtained from the relevant local ethical committees for medical research. All autopsies were performed within 10 hours after death. Paraffin-embedded tissues were sectioned at 6 _µ_m and mounted on pre-coated glass slides (StarFrost, Waldemar Knittel Glasbearbeitungs GmbH, Braunschweig, Germany). Four control patients without prior neurological disease and one ALS patient with a *FUS*-R521C mutation were analyzed. Spinal cord sections on the cervical, thoracic, and lumbar levels were processed as follows: At first, the sections were deparaffinized in xylene for 20 min and then rehydrated in 100%, 96%, and 70% ethanol for 5 min each followed by endogenous peroxidase quenching (0.3% H_2_O_2_ in methanol) for 20 min. Antigen retrieval was performed in these sections by heating them in citrate buffer, pH 6 (DAKO), for 20 min in a pressure cooker. After washing in PBS, sections were incubated with primary for 1 h at room temperature or 4°C overnight. After washing in PBS, sections were incubated with the appropriate secondary antibody (ImmunoLogic, Duiven, The Netherlands) for 30 min at room temperature. DAB reagent (ImmunoLogic, ready to use) was used to visualize antibody binding. The sections were then counter-stained with 6% hematoxylin for 3 min. All procedures were performed at room temperature.

### RNA-sequencing and analysis

RNA-seq anlalysis of somatodendritic (SD) and axonal compartments was performed as described in detail in Zimyanin V., Dash BP. (Reference). The generation of human NPCs and sMNs was accomplished following the modifications protocol from Reinhardt et al. (2013) and described also in Naumann, Pall, et al., 2018 ^51,52^. Briefly, cells were grown in Xona (RD900) microfluidic devices (7-10 per sample per phenotype) and total RNA isolated from either SD or axonal compartment after 2 weeks of growth. Total RNA was sequenced (at Biotech Deep sequencing facility, TU, Dresden) on an Illumina HiSeq 2500 sequencing system using Illumina sequencing libraries. An average of about 30 million 75 base pairs long single-end reads were produced for each sample for RNA-seq. The sequencing reads were aligned to the human hg38/GRCh38 reference genome and EnsEMBL assembly v81 gene annotation model using GSNAP aligner (v2017-11-15) (Wu and Nacu, 2010) with splice-junction support from annotated genes (EnsEMBL v81), respectively. A table of raw read counts per gene was obtained based on the overlap of the uniquely mapped reads with annotated human genes (EnsEMBL v81) using featureCounts (v1.5.2) (Liao et al., 2014). Data normalization and differential expression were analysed using the DESeq2 R package v1.16.1 (Love et al., 2014) with a false discovery rate (FDR) ≤ 0.05 as the significant cutoff. The PPI networks were performed with the Search Tool for the Retrieval of Interacting Genes (STRING) database v12.0 ^53^ and Cytoscape v3.10.1 ^54^. The datasets analysed during the current study are available at GSE276214.

## Institutional Review Board Statement

The performed procedures were in accordance with the Declaration of Helsinki (WMA, 1964) and approved by the Ethical Committee of the Technische Universität Dresden, Germany (EK 393122012 and EK 45022009) and Rostock (A2019-0137).

## Informed Consent Statement

Written informed consent was obtained from all participants including for publication of any research results.

## Author contributions according to CRediT

Conceptualization (MN, AH); Data curation (MN, SK, VZ); Formal Analysis (MN, MK, SK, VZ, PO, BPD); Funding acquisition (AH); Methodology (MN, DG, VZ, MK, SK, MLK, PO); Project administration (AH); Resources (MN, EA, KP, RG, VZ, MK, HG, FP, MLK, SK, DB, SP, TB, JS, AR, TG, PÖ, AH); Software (MN, HG, DG, VZ, BPD); Supervision (AH); Validation (MN, DG); Visualization (MN, AH); Writing – original draft (MN, AH); Writing – review & editing (all authors)

## Acknowledgments

We acknowledge the great cell culture help of Jette Abel and Sylvia Kanzler Additionally, we express our gratitude to Franziska Alfen for her Western Blot support. We thank Susann Lehmann for performing histological stains. We thank J.J. Anink (Amsterdam UMC neuropathology) for technical support. We thank Stephen Meier (Ulm) for excellent technical assistance with the Ella NfL measurements.

## Data availability

We have included all relevant data in the manuscript. The RNA-seq datasets were obtained from GEO database (Accession No. GSE276214) (Zimyanin V., Dash BP). More details are available from the authors.

---

**Supplement Figure 1:** (A) IF panel of FUS^wt^ and FUS-P525L sMNs indicating yH2A.x as a measure of DNA double-strand breaks, scale bar 10µM. Etoposide 2µM 72h was used as a positive control. Note the increased amount of yH2A.x nuclear foci in FUS^mut^ sMNs at baseline. (B) IF image panel of FUS^wt^ and FUS-P525L sMNs according to Fig. 2A. Cells were stained against the mitochondrial outer membrane protein TOM20 and the anti-dsRNA (J2) antibody. Nuclear staining was done with DAPI. C-terminally tagged FUS-GFP indicates FUS presence in the different conditions and is visualized via the Green-Fire-Blue LUT in Fiji. (C) WB scan for MDA5 and RIG-I in sMNs with either FUS^wt^ or FUS-P525L mutation. Treatment with IFN-beta 100ng/ml 24h as a positive control. On baseline, no signal for MDA5 was detected in either cell line for MDA5 in contrast to RIG-I. (D) WB scan for phospo-p65(p-RelA) in FUS^wt^ or FUS-P525L sMNs protein samples. TNF-alpha 20ng/ml for 24h was used as a positive control indicated by the slight increase in band intensity on the third and fourth lanes.

## References

1. Weishaupt, J.H., Hyman, T., and Dikic, I. (2016). Common Molecular Pathways in Amyotrophic Lateral Sclerosis and Frontotemporal Dementia. Trends Mol Med 22, 769–783. 10.1016/j.molmed.2016.07.005.

2. Gulen, M.F., Samson, N., Keller, A., Schwabenland, M., Liu, C., Gluck, S., Thacker, V.V., Favre, L., Mangeat, B., Kroese, L.J., et al. (2023). cGAS-STING drives ageing-related inflammation and neurodegeneration. Nature 620, 374–380. 10.1038/s41586-023-06373-1.

3. Sliter, D.A., Martinez, J., Hao, L., Chen, X., Sun, N., Fischer, T.D., Burman, J.L., Li, Y., Zhang, Z., Narendra, D.P., et al. (2018). Parkin and PINK1 mitigate STING-induced inflammation. Nature 561, 258–262. 10.1038/s41586-018-0448-9.

4. Yu, C.H., Davidson, S., Harapas, C.R., Hilton, J.B., Mlodzianoski, M.J., Laohamonthonkul, P., Louis, C., Low, R.R.J., Moecking, J., De Nardo, D., et al. (2020). TDP-43 Triggers Mitochondrial DNA Release via mPTP to Activate cGAS/STING in ALS. Cell 183, 636–649 e618. 10.1016/j.cell.2020.09.020.

5. Ablasser, A., Goldeck, M., Cavlar, T., Deimling, T., Witte, G., Rohl, I., Hopfner, K.P., Ludwig, J., and Hornung, V. (2013). cGAS produces a 2’-5’-linked cyclic dinucleotide second messenger that activates STING. Nature 498, 380–384. 10.1038/nature12306.

6. Wu, B., and Hur, S. (2015). How RIG-I like receptors activate MAVS. Curr Opin Virol 12, 91–98. 10.1016/j.coviro.2015.04.004.

7. Liu, S., Cai, X., Wu, J., Cong, Q., Chen, X., Li, T., Du, F., Ren, J., Wu, Y.T., Grishin, N.V., and Chen, Z.J. (2015). Phosphorylation of innate immune adaptor proteins MAVS, STING, and TRIF induces IRF3 activation. Science 347, aaa2630. 10.1126/science.aaa2630.

8. Pang, W., and Hu, F. (2023). C9ORF72 suppresses JAK-STAT mediated inflammation. iScience 26, 106579. 10.1016/j.isci.2023.106579.

9. Rodriguez, S., Sahin, A., Schrank, B.R., Al-Lawati, H., Costantino, I., Benz, E., Fard, D., Albers, A.D., Cao, L., Gomez, A.C., et al. (2021). Genome-encoded cytoplasmic double-stranded RNAs, found in C9ORF72 ALS-FTD brain, propagate neuronal loss. Sci Transl Med 13. 10.1126/scitranslmed.aaz4699.

10. McCauley, M.E., O’Rourke, J.G., Yanez, A., Markman, J.L., Ho, R., Wang, X., Chen, S., Lall, D., Jin, M., Muhammad, A., et al. (2020). C9orf72 in myeloid cells suppresses STING-induced inflammation. Nature 585, 96–101. 10.1038/s41586-020-2625-x.

11. Tan, H.Y., Yong, Y.K., Xue, Y.C., Liu, H., Furihata, T., Shankar, E.M., and Ng, C.S. (2022). cGAS and DDX41-STING mediated intrinsic immunity spreads intercellularly to promote neuroinflammation in SOD1 ALS model. iScience 25, 104404. 10.1016/j.isci.2022.104404.

12. Marques, C., Held, A., Dorfman, K., Sung, J., Song, C., Kavuturu, A.S., Aguilar, C., Russo, T., Oakley, D.H., Albers, M.W., et al. (2024). Neuronal STING activation in amyotrophic lateral sclerosis and frontotemporal dementia. Acta Neuropathol 147, 56. 10.1007/s00401-024-02688-z.

13. Naumann, M., Pal, A., Goswami, A., Lojewski, X., Japtok, J., Vehlow, A., Naujock, M., Gunther, R., Jin, M., Stanslowsky, N., et al. (2018). Impaired DNA damage response signaling by FUS-NLS mutations leads to neurodegeneration and FUS aggregate formation. Nature communications 9, 335. 10.1038/s41467-017-02299-1.

14. Zimyanin, V.L., Pielka, A.M., Glass, H., Japtok, J., Grossmann, D., Martin, M., Deussen, A., Szewczyk, B., Deppmann, C., Zunder, E., et al. (2023). Live Cell Imaging of ATP Levels Reveals Metabolic Compartmentalization within Motoneurons and Early Metabolic Changes in FUS ALS Motoneurons. Cells 12. 10.3390/cells12101352.

15. Guo, W., Naujock, M., Fumagalli, L., Vandoorne, T., Baatsen, P., Boon, R., Ordovas, L., Patel, A., Welters, M., Vanwelden, T., et al. (2017). HDAC6 inhibition reverses axonal transport defects in motor neurons derived from FUS-ALS patients. Nat Commun 8, 861. 10.1038/s41467-017-00911-y.

16. Guo, W., Wang, H., Kumar Tharkeshwar, A., Couthouis, J., Braems, E., Masrori, P., Van Schoor, E., Fan, Y., Ahuja, K., Moisse, M., et al. (2023). CRISPR/Cas9 screen in human iPSC-derived cortical neurons identifies NEK6 as a novel disease modifier of C9orf72 poly(PR) toxicity. Alzheimers Dement 19, 1245–1259. 10.1002/alz.12760.

17. De Vos, K., Severin, F., Van Herreweghe, F., Vancompernolle, K., Goossens, V., Hyman, A., and Grooten, J. (2000). Tumor necrosis factor induces hyperphosphorylation of kinesin light chain and inhibits kinesin-mediated transport of mitochondria. J Cell Biol 149, 1207–1214. 10.1083/jcb.149.6.1207.

18. Viengkhou, B., and Hofer, M.J. (2023). Breaking down the cellular responses to type I interferon neurotoxicity in the brain. Front Immunol 14, 1110593. 10.3389/fimmu.2023.1110593.

19. Rulten, S.L., Rotheray, A., Green, R.L., Grundy, G.J., Moore, D.A., Gomez-Herreros, F., Hafezparast, M., and Caldecott, K.W. (2014). PARP-1 dependent recruitment of the amyotrophic lateral sclerosis-associated protein FUS/TLS to sites of oxidative DNA damage. Nucleic Acids Res 42, 307–314. 10.1093/nar/gkt835.

20. Haag, S.M., Gulen, M.F., Reymond, L., Gibelin, A., Abrami, L., Decout, A., Heymann, M., van der Goot, F.G., Turcatti, G., Behrendt, R., and Ablasser, A. (2018). Targeting STING with covalent small-molecule inhibitors. Nature 559, 269–273. 10.1038/s41586-018-0287-8.

21. Dhir, A., Dhir, S., Borowski, L.S., Jimenez, L., Teitell, M., Rotig, A., Crow, Y.J., Rice, G.I., Duffy, D., Tamby, C., et al. (2018). Mitochondrial double-stranded RNA triggers antiviral signalling in humans. Nature 560, 238–242. 10.1038/s41586-018-0363-0.

22. Irwin, K.E., Sheth, U., Wong, P.C., and Gendron, T.F. (2024). Fluid biomarkers for amyotrophic lateral sclerosis: a review. Mol Neurodegener 19, 9. 10.1186/s13024-023-00685-6.

23. Zimyanin, V., Dash, B.P., Großmann, D., Simolka, T., Glaß, H., Verma, R., Khatri, V., Deppmann, C., Zunder, E., Redemann, S., and Hermann, A. (2024). Axonal transcriptome reveals upregulation of PLK1 as a protective mechanism in response to increased DNA damage in FUS^P525L^ spinal motor neurons. bioRxiv, 2024.2011.2020.624439. 10.1101/2024.11.20.624439.

24. Bonekamp, N.A., Peter, B., Hillen, H.S., Felser, A., Bergbrede, T., Choidas, A., Horn, M., Unger, A., Di Lucrezia, R., Atanassov, I., et al. (2020). Small-molecule inhibitors of human mitochondrial DNA transcription. Nature 588, 712–716. 10.1038/s41586-020-03048-z.

25. Arnaiz, E., Miar, A., Dias Junior, A.G., Prasad, N., Schulze, U., Waithe, D., Nathan, J.A., Rehwinkel, J., and Harris, A.L. (2021). Hypoxia Regulates Endogenous Double-Stranded RNA Production via Reduced Mitochondrial DNA Transcription. Front Oncol 11, 779739. 10.3389/fonc.2021.779739.

26. Baechler, E.C., Batliwalla, F.M., Karypis, G., Gaffney, P.M., Ortmann, W.A., Espe, K.J., Shark, K.B., Grande, W.J., Hughes, K.M., Kapur, V., et al. (2003). Interferon-inducible gene expression signature in peripheral blood cells of patients with severe lupus. Proc Natl Acad Sci U S A 100, 2610–2615. 10.1073/pnas.0337679100.

27. Dunker, W., Ye, X., Zhao, Y., Liu, L., Richardson, A., and Karijolich, J. (2021). TDP-43 prevents endogenous RNAs from triggering a lethal RIG-I-dependent interferon response. Cell Rep 35, 108976. 10.1016/j.celrep.2021.108976.

28. Paget, M., Cadena, C., Ahmad, S., Wang, H.T., Jordan, T.X., Kim, E., Koo, B., Lyons, S.M., Ivanov, P., tenOever, B., et al. (2023). Stress granules are shock absorbers that prevent excessive innate immune responses to dsRNA. Mol Cell 83, 1180–1196 e1188. 10.1016/j.molcel.2023.03.010.

29. Maharana, S., Kretschmer, S., Hunger, S., Yan, X., Kuster, D., Traikov, S., Zillinger, T., Gentzel, M., Elangovan, S., Dasgupta, P., et al. (2022). SAMHD1 controls innate immunity by regulating condensation of immunogenic self RNA. Mol Cell 82, 3712–3728 e3710. 10.1016/j.molcel.2022.08.031.

30. Kim, S.S., Sze, L., Liu, C., and Lam, K.P. (2019). The stress granule protein G3BP1 binds viral dsRNA and RIG-I to enhance interferon-beta response. J Biol Chem 294, 6430–6438. 10.1074/jbc.RA118.005868.

31. Manivannan, P., Siddiqui, M.A., and Malathi, K. (2020). RNase L Amplifies Interferon Signaling by Inducing Protein Kinase R-Mediated Antiviral Stress Granules. J Virol 94. 10.1128/JVI.00205-20.

32. Szewczyk, B., Gunther, R., Japtok, J., Frech, M.J., Naumann, M., Lee, H.O., and Hermann, A. (2023). FUS ALS neurons activate major stress pathways and reduce translation as an early protective mechanism against neurodegeneration. Cell Rep 42, 112025. 10.1016/j.celrep.2023.112025.

33. Kodavati, M., Wang, H., Guo, W., Mitra, J., Hegde, P.M., Provasek, V., Rao, V.H.M., Vedula, I., Zhang, A., Mitra, S., et al. (2024). FUS unveiled in mitochondrial DNA repair and targeted ligase-1 expression rescues repair-defects in FUS-linked motor neuron disease. Nat Commun 15, 2156. 10.1038/s41467-024-45978-6.

34. Deng, J., Yang, M., Chen, Y., Chen, X., Liu, J., Sun, S., Cheng, H., Li, Y., Bigio, E.H., Mesulam, M., et al. (2015). FUS Interacts with HSP60 to Promote Mitochondrial Damage. PLoS genetics 11, e1005357. 10.1371/journal.pgen.1005357.

35. MacNair, L., Xiao, S., Miletic, D., Ghani, M., Julien, J.P., Keith, J., Zinman, L., Rogaeva, E., and Robertson, J. (2016). MTHFSD and DDX58 are novel RNA-binding proteins abnormally regulated in amyotrophic lateral sclerosis. Brain 139, 86–100. 10.1093/brain/awv308.

36. Honda, H., Yagita, K., Arahata, H., Hamasaki, H., Noguchi, H., Koyama, S., and Sasagasako, N. (2024). Increased expression of human antiviral protein MxA in FUS proteinopathy in amyotrophic lateral sclerosis. Brain Pathol 34, e13191. 10.1111/bpa.13191.

37. Ma, F., Li, B., Yu, Y., Iyer, S.S., Sun, M., and Cheng, G. (2015). Positive feedback regulation of type I interferon by the interferon-stimulated gene STING. EMBO Rep 16, 202–212. 10.15252/embr.201439366.

38. Zevini, A., Olagnier, D., and Hiscott, J. (2017). Crosstalk between Cytoplasmic RIG-I and STING Sensing Pathways. Trends Immunol 38, 194–205. 10.1016/j.it.2016.12.004.

39. Buss, H., Dorrie, A., Schmitz, M.L., Hoffmann, E., Resch, K., and Kracht, M. (2004). Constitutive and interleukin-1-inducible phosphorylation of p65 NF-kappaB at serine 536 is mediated by multiple protein kinases including IkappaB kinase (IKK)-alpha, IKKbeta, IKKepsilon, TRAF family member-associated (TANK)-binding kinase 1 (TBK1), and an unknown kinase and couples p65 to TATA-binding protein-associated factor II31-mediated interleukin-8 transcription. J Biol Chem 279, 55633–55643. 10.1074/jbc.M409825200.

40. Dunphy, G., Flannery, S.M., Almine, J.F., Connolly, D.J., Paulus, C., Jonsson, K.L., Jakobsen, M.R., Nevels, M.M., Bowie, A.G., and Unterholzner, L. (2018). Non-canonical Activation of the DNA Sensing Adaptor STING by ATM and IFI16 Mediates NF-kappaB Signaling after Nuclear DNA Damage. Mol Cell 71, 745–760 e745. 10.1016/j.molcel.2018.07.034.

41. Trishna, S., Lavon, A., Shteinfer-Kuzmine, A., Dafa-Berger, A., and Shoshan-Barmatz, V. (2023). Overexpression of the mitochondrial anti-viral signaling protein, MAVS, in cancers is associated with cell survival and inflammation. Mol Ther Nucleic Acids 33, 713–732. 10.1016/j.omtn.2023.07.008.

42. Xu, H., He, X., Zheng, H., Huang, L.J., Hou, F., Yu, Z., de la Cruz, M.J., Borkowski, B., Zhang, X., Chen, Z.J., and Jiang, Q.X. (2014). Structural basis for the prion-like MAVS filaments in antiviral innate immunity. Elife 3, e01489. 10.7554/eLife.01489.

43. Chattopadhyay, S., and Sen, G.C. (2017). RIG-I-like receptor-induced IRF3 mediated pathway of apoptosis (RIPA): a new antiviral pathway. Protein Cell 8, 165–168. 10.1007/s13238-016-0334-x.

44. Wang, H., Guo, W., Mitra, J., Hegde, P.M., Vandoorne, T., Eckelmann, B.J., Mitra, S., Tomkinson, A.E., Van Den Bosch, L., and Hegde, M.L. (2018). Mutant FUS causes DNA ligation defects to inhibit oxidative damage repair in Amyotrophic Lateral Sclerosis. Nat Commun 9, 3683. 10.1038/s41467-018-06111-6.

45. Deng, Q., Holler, C.J., Taylor, G., Hudson, K.F., Watkins, W., Gearing, M., Ito, D., Murray, M.E., Dickson, D.W., Seyfried, N.T., and Kukar, T. (2014). FUS is phosphorylated by DNA-PK and accumulates in the cytoplasm after DNA damage. J. Neurosci. 34, 7802–7813.

46. Gupta, P.K., Prabhakar, S., Sharma, S., and Anand, A. (2011). Vascular endothelial growth factor-A (VEGF-A) and chemokine ligand-2 (CCL2) in amyotrophic lateral sclerosis (ALS) patients. J Neuroinflammation 8, 47. 10.1186/1742-2094-8-47.

47. Irwin, K.E., Jasin, P., Braunstein, K.E., Sinha, I.R., Garret, M.A., Bowden, K.D., Chang, K., Troncoso, J.C., Moghekar, A., Oh, E.S., et al. (2024). A fluid biomarker reveals loss of TDP-43 splicing repression in presymptomatic ALS-FTD. Nat Med 30, 382–393. 10.1038/s41591-023-02788-5.

48. Chatterjee, M., Ozdemir, S., Fritz, C., Mobius, W., Kleineidam, L., Mandelkow, E., Biernat, J., Dogdu, C., Peters, O., Cosma, N.C., et al. (2024). Plasma extracellular vesicle tau and TDP-43 as diagnostic biomarkers in FTD and ALS. Nat Med 30, 1771–1783. 10.1038/s41591-024-02937-4.

49. Reinhardt, P., Glatza, M., Hemmer, K., Tsytsyura, Y., Thiel, C.S., Hoing, S., Moritz, S., Parga, J.A., Wagner, L., Bruder, J.M., et al. (2013). Derivation and expansion using only small molecules of human neural progenitors for neurodegenerative disease modeling. PLoS ONE 8, e59252–e59252.

50. Wolf, C., Bruck, N., Koss, S., Griep, C., Kirschfink, M., Palm-Beden, K., Fang, M., Rober, N., Winkler, S., Berner, R., et al. (2020). Janus kinase inhibition in complement component 1 deficiency. J Allergy Clin Immunol 146, 1439–1442 e1435. 10.1016/j.jaci.2020.04.002.

51. Naumann, M., Pal, A., Goswami, A., Lojewski, X., Japtok, J., Vehlow, A., Naujock, M., Günther, R., Jin, M., Stanslowsky, N., et al. (2018). Impaired DNA damage response signaling by FUS-NLS mutations leads to neurodegeneration and FUS aggregate formation. Nat Commun 9, 335. 10.1038/s41467-017-02299-1.

52. Reinhardt, P., Glatza, M., Hemmer, K., Tsytsyura, Y., Thiel, C.S., Hoing, S., Moritz, S., Parga, J.A., Wagner, L., Bruder, J.M., et al. (2013). Derivation and expansion using only small molecules of human neural progenitors for neurodegenerative disease modeling. PloS one 8, e59252. 10.1371/journal.pone.0059252.

53. Szklarczyk, D., Franceschini, A., Wyder, S., Forslund, K., Heller, D., Huerta-Cepas, J., Simonovic, M., Roth, A., Santos, A., Tsafou, K.P., et al. (2015). STRING v10: protein-protein interaction networks, integrated over the tree of life. Nucleic Acids Res 43, D447–452. 10.1093/nar/gku1003.

54. Shannon, P., Markiel, A., Ozier, O., Baliga, N.S., Wang, J.T., Ramage, D., Amin, N., Schwikowski, B., and Ideker, T. (2003). Cytoscape: a software environment for integrated models of biomolecular interaction networks. Genome Res 13, 2498–2504. 10.1101/gr.1239303.

