## Supplementary figures and images for "“RIG-I mediated neuron-specific IFN type 1 signaling in FUS-ALS induces neurodegeneration and offers new biomarker-driven individualized treatment options for (FUS-)ALS.”"

### Supplemental figure 1

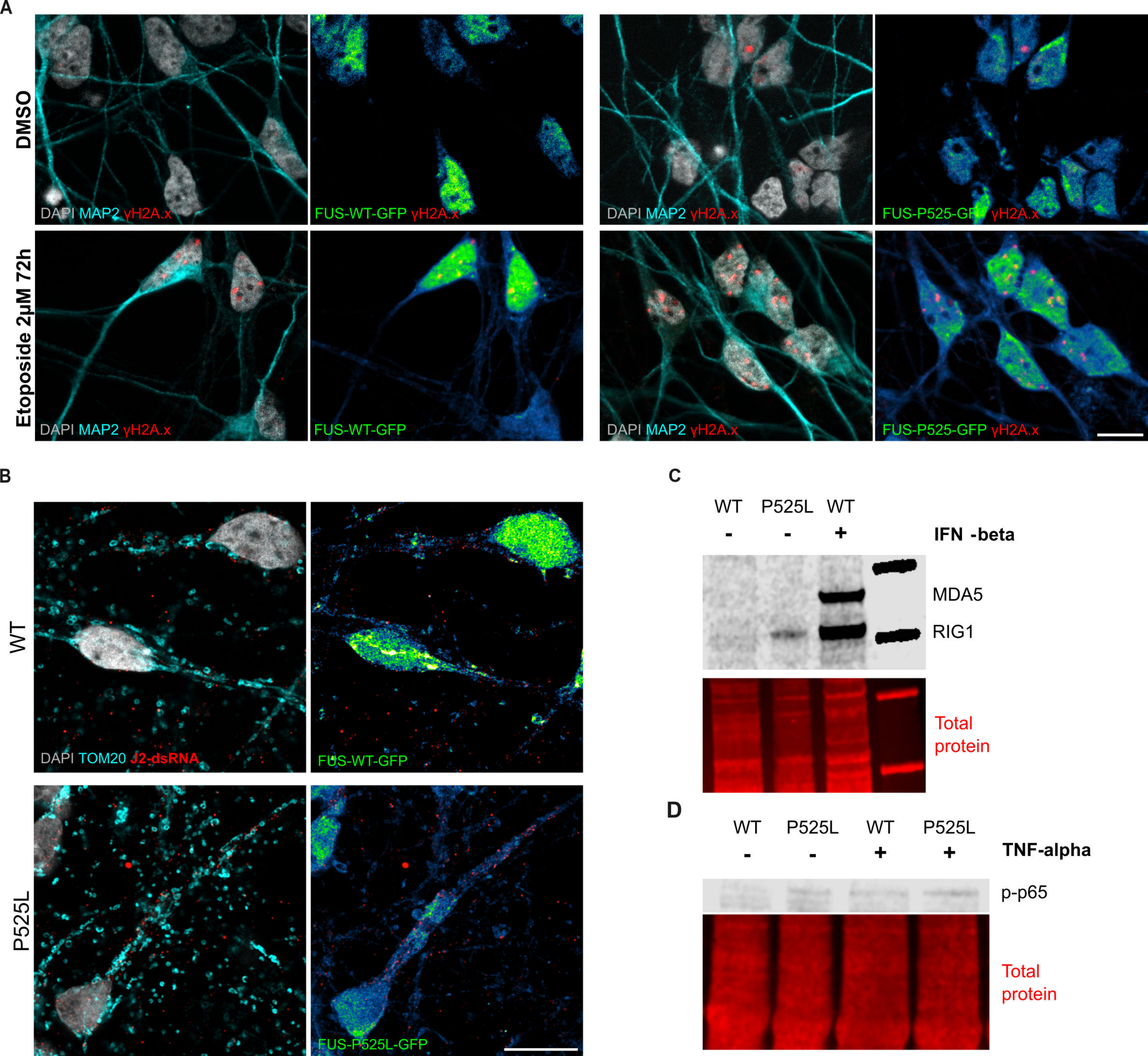
